# An ERF transcription factor from Brassica oleracea: a new member of the emerging pathogenicity hub in plant-Xanthomonas interactions

**DOI:** 10.1101/2020.08.21.259085

**Authors:** Nikolay Zlobin, Marina Lebedeva, Yuliya Monakhova, Vera Ustinova, Vasiliy Taranov

## Abstract

- TAL effectors (TALEs), which induce the expression of specific plant genes to promote infection, are the main pathogenic determinants of different Xanthomonas bacteria. However, investigation of TALEs from *Xanthomonas campestris* pv. *campestris*, which causes black rot disease of crucifers, is in its infancy.
- In this study, we used PCR-based amplification in conjunction with SMRT amplicon sequencing to identify TALE genes in several *Xanthomonas campestris* pv. *campestris* strains and performed computational prediction in conjunction with RT-PCR-based analysis to identify their target genes in *Brassica oleracea*.
- Transcription factor from the AP2/ERF family was predicted to be putative target gene for the conserved TALEs present in multiple *Xanthomonas campestris* pv. *campestris* strains. Its expression dramatically increased upon leaf inoculation with strains harbouring such TALEs.
- Several members of the AP2/ERF factor family from different plant species were identified as targets of TALEs from various Xanthomonas species, which suggests that they constitute a new pathogenicity hub in plant-Xanthomonas interactions.

## Introduction

Transcription activator-like effectors (TALEs) of Xanthomonas bacteria are trans-kingdom transcription factors that are translocated to plant cells via the type III secretion system and that specifically bind to the promoters of certain host genes and activate their transcription (Schandry *et al*., 2018). The central part of TALEs consist of a series of quasi-identical repeats typically comprising 33-35 amino acid (aa) residues, which differ mainly by residues at positions 12-13, known as repeat variable di-residues (RVDs). Each TALE repeat binds a single nucleotide, and repeats with different RVDs preferentially bind different nucleotides (Schandry *et al*., 2018). The order of repeats in a TALE determines the sequence in the plant genomic DNA to which the TALE specifically binds. If this sequence, which referred to as the effector-binding element (EBE), is located within the promoter region of a gene, TALEs activate the expression of the gene upon binding (Hutin *et al*., 2015; Schandry *et al*., 2018). Hence, if the primary structure of a TALE and, accordingly, the order of the RVD are known, it is possible to predict TALE target genes in a plant genome (Pérez-Quintero & Szurek, 2019).

TALE-mediated upregulation of target genes promotes disease development. TALE target genes are also called susceptibility genes, or S genes (Schandry *et al*., 2018). TALE-dependent induction of the expression of a single S gene can determine the difference between resistance and susceptibility in plant-Xanthomonas interactions (Verdier *et al*., 2012; Streubel *et al*., 2013; Hutin *et al*., 2015; Tran *et al*., 2018), and TALEs that activate such genes are often conserved and widely distributed among Xanthomonas species (Hutin *et al*., 2015; Oliva *et al*., 2019). Some TALE-activated S genes are conserved between different plant-Xanthomonas systems. The most well-studied examples are members of the SWEET gene family, which TALE-dependent activation was observed in the different plants (rice, citrus, cassava, cotton, pepper) upon infection with the different Xanthomonas species and strains (Hutin *et al*., 2015; Pérez-Quintero *et al*., 2015; Pérez-Quintero & Szurek, 2019). In some cases, this activation has been shown to be critically important for disease development (Hutin *et al*., 2015; Pérez-Quintero & Szurek, 2019). It is generally believed that SWEET overexpression increases sugar efflux from plant cells and facilitates bacterial colonization of the apoplast (Streubel *et al*., 2013). Another example of conserved TALE targets involve transcription factors. For example, different TALEs from several pathovars of *Xanthomonas citri* activate the expression of the LOB1 transcription factor in citrus, and TALEs from multiple *Xanthomonas oryzae* pv. *oryzae* strains upregulate the expression of TFX1 transcription factors (Sugio *et al*., 2007; Hutin *et al*., 2015; Pérez-Quintero *et al*., 2015; Pérez-Quintero & Szurek, 2019). Conservative S genes or S gene families are often referred to as susceptibility hubs or pathogenicity hubs (Hutin *et al*., 2015; Mücke *et al*., 2019; Pérez-Quintero & Szurek, 2019; Wu *et al*., 2019). Their identification is important not only for improving our understanding of the infection process but also for developing resistant plants, because the modification of conserved TALE targets can lead to more durable resistance.

*Xanthomonas campestris* pv. *campestris* (Xcc) causes black rot, the most harmful and economically important disease of vegetable Brassica crops (Vicente *et al*., 2013). The presence of TALEs has been shown for many Xcc strains of different geographical origins (Mokryakov *et al*., 2010; Denancé *et al*., 2018). However, studies of the role of Xcc TALEs in black rot disease development are in their infancy, and TALE target genes and the means by which their activation promotes the infection process are largely unknown. In this study, the *Brassica oleracea* target gene of Xcc TALEs with a common RVD combination was discovered. This gene belongs to a transcription factor gene family and represents a new member of the emerging susceptibility hub.

## Materials and methods

### Bacterial strains, growth conditions and Xcc genomic DNA extraction

The following Xcc strains of different geographical origins were used in this study: DK-1, Ram 1-3, Ram 2-2, Ram 3-2, Ram 4-1, B-1, and Bun-1 (from the Moscow region); Tir2, XY1-1, XY1-2, XY2-1, and XY2-2 (from Crimea); 306NZ and 276NZ (from The Netherlands); and Xn-13 (from Japan). The strains are referred to Xcc races 1, 3, 4, and 6 according to the report of Ha *et al*. (2014). For genomic DNA extraction, Xcc was cultured on solid LB media. Genomic DNA extraction was performed using the cetyl-trimethylammonium bromide (CTAB) method.

### TALE gene amplification and sequencing

Amplification of TALE genes was performed via a two-step PCR process in conjunction with primers 5’-GATCCCATTCGTTCGCGCACACCAAGTC-3’ and 5’-CTCCATCAACCATGCGAGCTCCTCTTCG-3’ and Taq DNA polymerase (Evrogen, Russia) under the following conditions: initial denaturation at 95°C for 4 min, 32 cycles of 95°C for 0.5 min followed by 75°C for 2.5 min, and a final elongation at 75°C for 5 min using the TP4-PCR-01 “Tertsik” thermocycler (DNA Technology, Russia).

The PCR products were visualized on a 2% agarose gel (TopVision Agarose, Thermo Scientific, USA) and extracted using a GeneJET Gel Extraction Kit (Thermo Fisher Scientific, USA). The full-length TALEs were sequenced using a MinION sequencer (Oxford Nanopore Technologies, UK), and libraries were prepared using a SQK-LSK109 ligation sequencing kit together with barcodes from an EXP-PBC001 Barcoding Kit (Oxford Nanopore Technologies, UK). The library contents were loaded onto a single MinION R9.4.1 flow cell and sequenced for 24 hrs.

Base calling and subsequent analysis of the reads were performed using Guppy 2.1.3 software (Oxford Nanopore Technologies). The reads were aligned using the ClustalW algorithm within UniPro UGENE software (Okonechnikov *et al*., 2012), and consensus sequences for the TALEs from each strain of the 6^th^ Xcc race were determined. In all 4 TALE genes, the region harbouring several consecutive cytosines was within the fifth repeat, but the exact number could not be determined based on the single-molecule real-time (SMRT) data. The sequence of the fifth repeat was determined using Sanger sequencing.

### Phylogenetic analyses

Phylogenetic analysis of the Xcc TALEs was performed using the DisTAL program (Pérez-Quintero *et al*., 2015). The analysis involved 4 TALE genes sequenced in this work and previously identified full-length TALE genes (Denancé *et al*., 2018). TALEs from *Xanthomonas translucens* pv. *undulosa* were used as the outgroup. All TALE sequences were aligned manually according to the instructions, and a phylogenetic tree was constructed without considering RVD sequences in the repeats.

### EBE prediction

TALE target prediction was performed using the PrediTALE tool (Erkes *et al*., 2019) for the genome of the *B. oleracea* TO1000DH3 line (NCBI Assembly: GCA_000695525.1) (Parkin *et al*., 2014). The default parameters were used, but the minus-strand EBE location penalty was omitted. Nine targets were associated with the maximal score (≈0.68), while the other targets had scores lower than 0.64. Genome sequences around these 9 EBEs were searched manually to identify annotated genes via the BLAST algorithm of the NCBI database (Sayers *et al*., 2020).

### Plant material, growth conditions and leaf inoculation

The *B. oleracea* kale-like homozygous double-haploid line TO1000DH3 was obtained from the Arabidopsis Biological Resource Center (stock number: CS29002) using the Arabidopsis Information Resource (TAIR) database (Berardini *et al*., 2015). The plants were grown in a moss peat:perlite mixture (10:1) watered with 10x diluted MS medium (Duchefa Biochemie, The Netherlands). Two Xcc strains belonging to race 6 were used for inoculation: XY1-1 and XY2-1. For inoculation, Xcc was cultivated on solid YDC media (Goszczynska *et al*., 2000). Fully expanded leaves of 8-week-old plants were used for inoculation with a 1 ml needleless insulin syringe, with bacteria resuspended to an OD_600_ of 0.1 (approximately 3×10^7^ CFU/ml) in 10 mm MgCl_2_. Three to four leaves per plant were inoculated.

### TALE target validation assays by qRT-PCR

The inoculated areas of leaves were excised at 6, 12, 24, and 48 hrs post-inoculation (hpi). Four fragments from different inoculated leaves were mixed together into a single sample to isolate the total RNA using TRIzol reagent (Thermo Fisher Scientific). cDNA was obtained using a ProtoScript II First Strand cDNA Synthesis Kit (NEB, USA) according to the manufacturer’s instructions. Two samples (biological replicates) were obtained for each strain + hpi combination. RT-PCR was then performed using a CFX-96 Real-Time PCR Detection System (Bio-Rad, USA) in conjunction with an M-427 ready qRT-PCR mixture (Syntol, Russia) and primers (0.2 μM) under the following conditions: initial denaturation at 95°C for 5 min, 45 cycles of 65°C for 0.5 min followed by 95°C for 0.25 min, and then a melting from 65°C to 95°C at a rate of a 0.5°C decrease every 5 seconds. The primers used for qRT-PCR are listed in Supporting Information Table S1. The specificity of amplification for each primer pair was verified by melting curve analysis. Three reference genes were used to normalize the expression. Amplification efficiency was calculated via a serial dilution of cDNA. The expression data were analysed using qbase+ software (Biogazelle, Belgium). In the resulting graph, each bar represents the average of two biological replicates (with three technical replicates each) together with the standard error of the mean. Once the relative expression in units was calculated, the expression in the mock sample at 48 hpi was considered one relative unit.

## Results

### Xcc strains of the 6 race contain the single TALE with a common RVD combination

The use of amplification-based methods to study TALE genes is challenging due to their large size, high GC content and presence of an array of quasi-identical repeats (Morbitzer *et al*., 2010; Hommelsheim *et al*., 2014; Pérez-Quintero & Szurek, 2019). To isolate the TALE genes, we used two-step high-temperature PCR with primers that anneal far from the repeat region, which greatly facilitates amplification according to Hommelsheim *et al*. (2014). In the sample of Xcc strains from the different races, all 4 strains belonging to Xcc race 6 carried a single TALE gene, whereas strains from races 1, 3 and 4 carried none (Fig. 1). Since the primers used for amplification anneal at the conserved regions of the TALE genes, they apparently can be used for the inexpensive and rapid identification and isolation of TALE genes from diverse Xanthomonas species.

**Figure 1.**
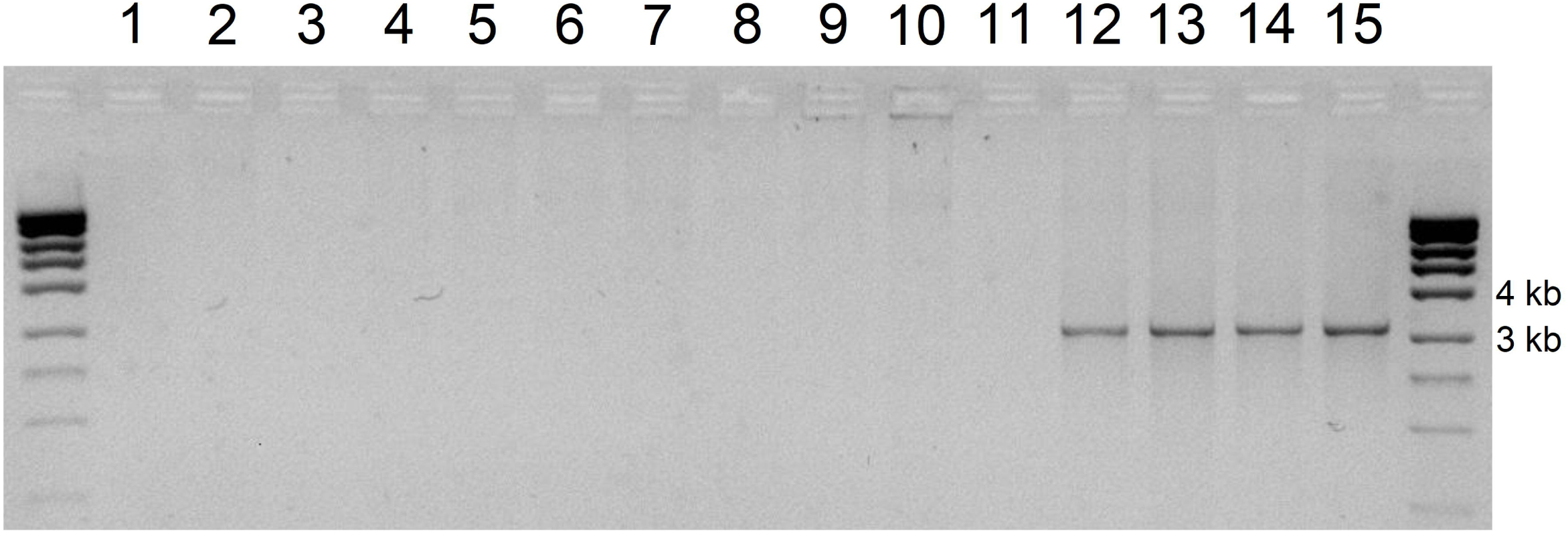
TALE gene identification in the following Xcc strains: DK-1 (1), Ram 3-2 (2), Ram 4-1 (3), 276NZ (4), and Tir-2 (5) – race 1; Ram 1-3 (6), Ram 2-2 (7), 306NZ (8), and B-1 (9) – race 3; Bun-1 (10) and Xn-13 (11) – race 4; XY1-1 (12), XY1-2 (13), XY2-1 (14), and XY2-2 (15) – race 6. Marker – MassRuler High-Range DNA ladder (Thermo Fisher Scientific).

SMRT amplicon sequencing showed that the TALE genes from all 4 strains contained 15 repeats (14 full repeats, with the last one truncated). According to the TALE Class Assignment algorithm from the AnnoTale suite (Grau *et al*., 2016), they were assigned to the TalEN class and named TalEN5 (Accession Number MT828881, strain XY1-1), TalEN6 (MT828882, strain XY1-2), TalEN7 (MT828883, strain XY2-1), and TalEN8 (MT828884, strain XY2-2). Phylogenetic relationships between TalEN(5-8) and previously sequenced Xcc TALEs (Denancé *et al*., 2018) were studied using the DisTAL algorithm (Pérez-Quintero *et al*., 2015). On the phylogenetic tree, TalEN(5-8) were located closest to TALEs from the Chinese strains CN-12, CN-17, and CN-18 (Fig. 2a). These TALEs also belong to the TalEN class according to the AnnoTale suite and were referred to as members of the Tal15g group by Denancé *et al*. (2018).

**Figure 2.**
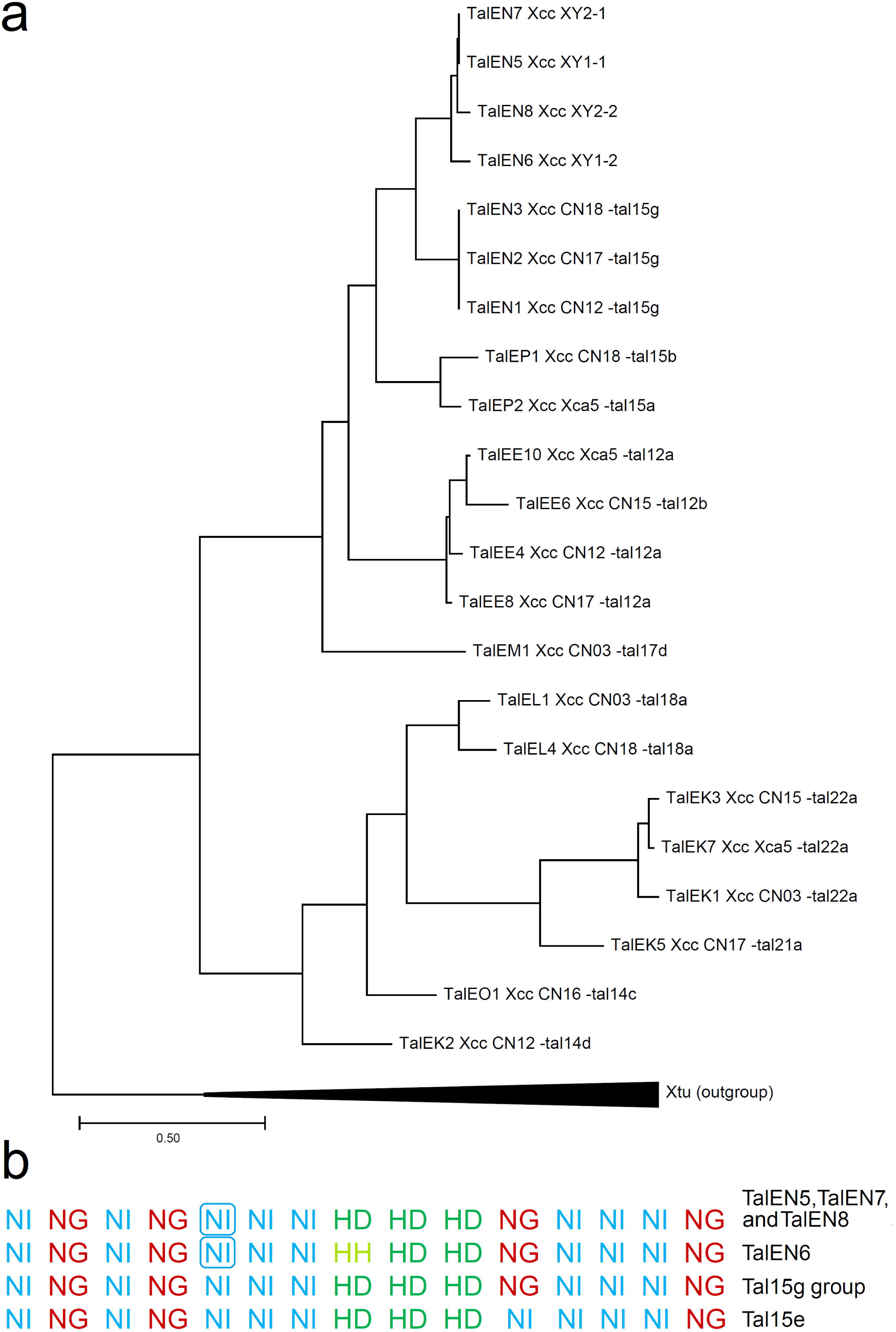
**A**. Phylogenetic tree of the full-length TALE genes sequenced in this study and in the study by Denancé *et al*. (2018). For each TALE gene from Denancé et al. (2018), corresponding RVD group name is provided. **B**. RVD sequences of TALEs from this study and the Tal15 g group and Tal15e from the study by Denancé *et al*. (2018). The RVDs from the fifth repeats of TalEN(5-8), which are 35 aas in length, are framed.

Differences in the nucleotide sequences between TalEN(5-8) and Tal15g group members occurred mainly at the 3’-end of the coding region (Supporting Information Figure S1). Additionally, all repeats in the central part of the Tal15g TALEs were 102 nucleotides in length and encoded 34 aa protein repeats, while in TalEN(5-8), the fifth repeat was 105 nucleotides in length and coded for a 35 aa repeat. It was shown previously that in different Xcc TALEs, all 35 aa repeats end with the «PHD/C» motif, whereas 34 aa repeats end with the «HG» motif (Denancé *et al*., 2018), and the same pattern was observed for the repeats in TalEN(5-8). The combination of repeats of different lengths in a single protein is generally not very common for TALEs from different Xanthomonas species but is typical for Xcc TALEs (Denancé *et al*., 2018).

Despite some differences in both the nucleotide and aa sequences between TalEN(5-8) and TALEs from the Tal15g group, the composition and order of RVDs were identical between them (Fig. 2b). The only exception was TalEN6 from the XY1-2 strain, which contains an HH RVD instead of an HD RVD in repeat 8; however, repeats with both HH and HD RVDs preferentially bind cytosine in the genomic DNA. The geographical origin of strains harbouring such RVD combinations varies – China and Belgium according to the work of Denancé *et al*. (2018) and Crimea in this work. TALEs from the Tal15g group were found in 7 of the 22 Xcc strains, which harbour any TALE genes according to Denancé *et al*. (2018). Also very similar RVD organization had Tal15e (Fig. 2b) (Denancé *et al*., 2018). This suggests that such RVD combination is common among Xcc TALEs. Identification of a larger number of Xcc strains of different geographical origins is desirable to determine whether such RVD combination arise independently. Since the RVD composition defines a set of TALE target genes, the existence of identical RVD arrays in TALEs from multiple Xcc strains of different geographical origins suggests the importance of their targets as S genes upon infection (Mücke *et al*., 2019).

### The ERF121 gene of *B. oleracea* is the putative target for TalEN(5-8) and is activated upon Xcc inoculation

The PrediTALE tool (Erkes *et al*., 2019) was used to identify putative EBEs for TalEN(5-8) in the genome of the *B. oleracea* TO1000DH3 line, which is highly susceptible to Xcc strains XY1-1, XY1-2, XY2-1, and XY2-2 upon vein inoculation (data not shown). One EBE received the highest score and was located in the promoter region of the annotated *B*.*oleracea* gene (GenBank Number XM_013739306.1), which codes for the putative ethylene-responsive transcription factor ERF121. The nucleotide sequence of this EBE is optimal for TalEN(5-8) binding (Fig. 3a). This is quite unusual because (with very rare exceptions, Mücke *et al*. (2019)) all known natural EBEs, including those from highly induced genes, harbour single or even multiple mismatches relative to the optimal TALE binding sequence. Since even a single mismatch can significantly reduce the extent of TALE-dependent gene activation (Erkes *et al*., 2017; Zaka *et al*., 2018; Cohn *et al*., 2016), the perfect match between the TalEN(5-8) RVD array and the EBE in the ERF121 promoter may be especially suitable for gene activation. In addition to the optimal sequence, the placement of EBE in the ERF121 promoter is also favourable. The EBE is located upstream of the CDS close to the putative start codon (Fig. 3a), which is typical for TALE-activated genes (Grau *et al*., 2013; Pereira *et al*., 2014; Wang *et al*., 2017; Streubel *et al*., 2017). Additionally, TALE EBEs often overlap with TATA boxes (Grau *et al*., 2013; Pereira *et al*., 2014; Cohn *et al*., 2016), and the EBE in ERF121 promoter comprises a TATA-like sequence (TATAWA consensus; Bernard *et al*. (2010)) (Fig. 3a).

**Figure 3.**
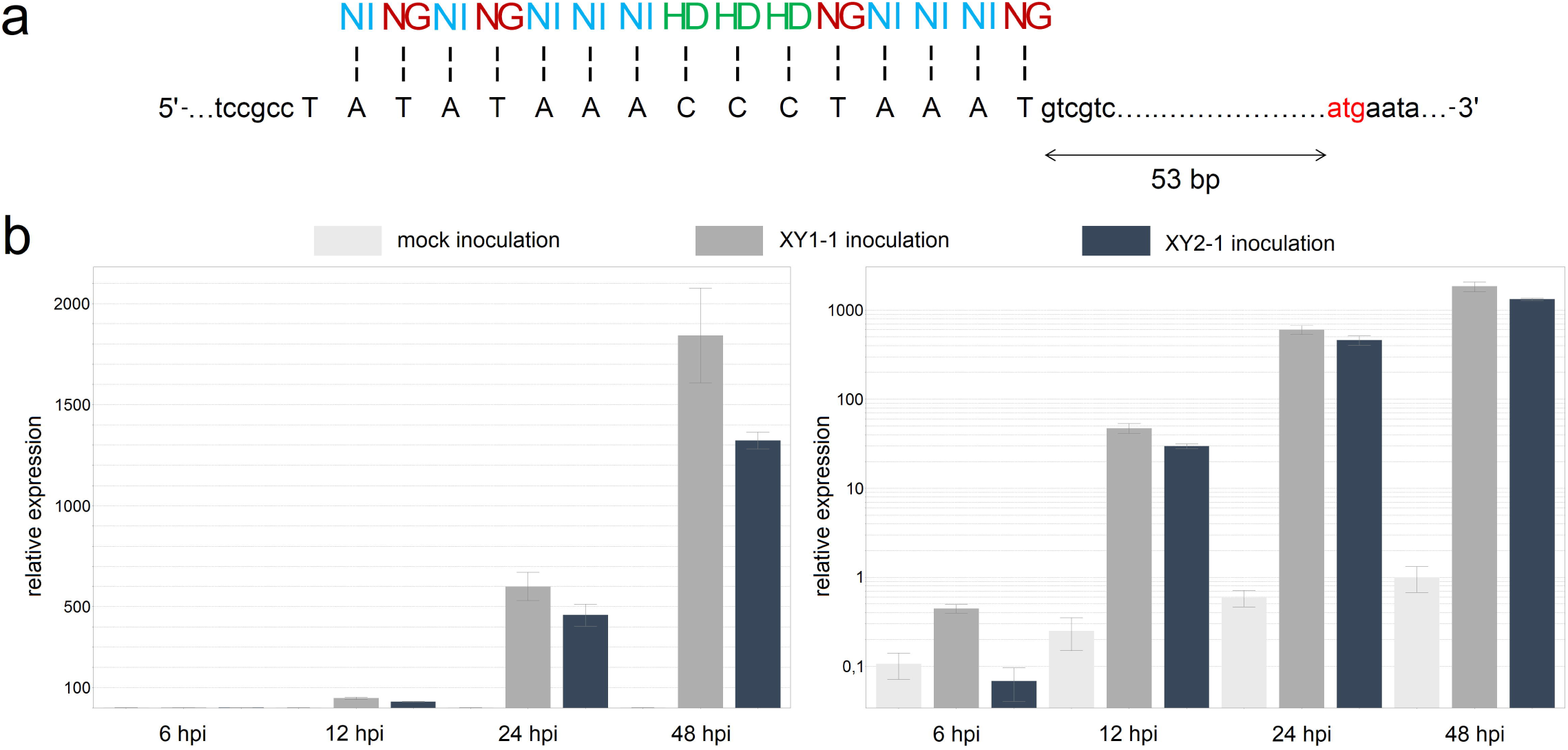
**A**. Composition and location of the EBE in the ERF121 gene promoter. The vertical dashes indicate matches between the EBE and RVDs. The putative start codon is highlighted in red. **B**. Expression level of ERF121 upon leaf inoculation at 6, 12, 24, and 48 hpi. Left – linear scale, right – logarithmic scale. Expression in the sample after mock inoculation at 48 hpi is considered a single expression unit. The bars show relative expression fold changes ±SEMs.

Due to the presence of the optimal EBE, ERF121 was considered a target for TalEN(5-8), and the expression of ERF121 was studied in *B. oleracea* upon inoculation with XY1-1 (carrying TalEN5) and XY2-1 (carrying TalEN7) strains. A substantial increase in ERF121 expression was observed at 12 hpi in response to both Xcc strains, and the expression increased several hundred times at 24 hpi and more than a thousand times at 48 hpi in comparison with that in response to the mock inoculation (Fig. 3b). Interestingly, ERF121 induction after inoculation with XY1-1 was observed to have occurred earlier, with a modest increase already at 6 hpi, and the expression generally was slightly higher than that after inoculation with XY2-1. Although TalEN5 and TalEN7 have identical RVDs, some influence on TALE activity may have non-RVD residues (Morbitzer *et al*., 2010), and also the effectivity of TALE synthesis (Hummel *et al*., 2017) and/or transport from the bacterial cell, which may differ between strains XY1-1 and XY2-1. We also observed an increase in ERF121 expression at different time points in response to the mock inoculation (Fig. 3b), which is not unusual in such studies (Sugio *et al*., 2007; Peng *et al*., 2019).

ERF121 belongs to the large ERF group within the AP2/ERF transcription factor family (Licausi *et al*., 2013). Several members of this family from diverse plant species have been previously identified as TALE targets (Pérez-Quintero *et al*., 2013; Wang *et al*., 2017; Tran *et al*., 2018; Peng *et al*., 2019), but they have limited homology to *Brassica oleracea* ERF121 or to each other and fall into the different phylogenetic groups within the AP2/ERF family (Supporting Information Table S2), which suggests dissimilar functions. Numerous ERF transcription factors have been shown to be involved in the plant response to various stress factors (Gutterson & Reuber, 2004; Licausi *et al*., 2013). Although the expression of ERF121 relatives from *A. thaliana* (AT2G20350, AT5G67000, AT5G67010) changed under stress conditions and treatment with some defense-associated plant hormones (Feng *et al*., 2005; McGrath *et al*., 2005; Postnikova *et al*., 2011; Pierce & Rey, 2013; Caarls *et al*., 2017), direct evidence of the involvement of *B. oleracea* ERF121 in plant-pathogen interactions are lacking.

## Discussion

Because of their role as the main pathogenic determinants, TALEs have been widely studied in Xanthomonas pathogens of several crop species, especially rice, citrus and pepper. In sharp contrast, the role of TALEs in Xanthomonas interactions with Brassicaceae family members, which include several important crop species, has surprisingly received little attention, with only a few studies having been published (Kay *et al*., 2005; Denancé *et al*., 2018). In this work, in *B. oleracea*, we identified the target gene for Xcc TALEs with conserved RVD array. The presence of the optimal EBE and the strong induction already detected at 12-24 hpi indicate that ERF121 is a direct target, while the presence of TALEs with such RVD combination in multiple Xcc strains of different origin suggests that ERF121 activation is a widespread Xcc pathogenic strategy.

Transcription factors are highly promising TALE targets because activation of a single such gene can lead to downstream changes in the expression of numerous genes and global shifts in the cellular environment. TALE-dependent activation of AP2/ERF transcription factors occurs in different plant species, such as rice (Pérez-Quintero *et al*., 2013; Wang *et al*., 2017; Tran *et al*., 2018; Tariq *et al*., 2019), wheat (Peng *et al*., 2019) and cabbage (this work), upon infection with various Xanthomonas species and strains. For example, expression of the ERF#123 gene was activated in rice by different TALEs from *Xanthomonas oryzae* pv. *oryzae* and *Xanthomonas oryzae* pv. *oryzicola* strains, and this activation increased rice susceptibility (Tran *et al*., 2018). ERF upregulation was also characteristic of the susceptible interaction between common bean and *Xanthomonas phaseoli* pv. *phaseoli* (Foucher *et al*., 2020). Interestingly, type III effector-dependent activation of many AP2/ERF transcription factors was observed in *Arabidopsis thaliana* upon *Pseudomonas syringae* pv. *tomato* infection (Truman *et al*., 2006). Taking into account the aforementioned experimental data, AP2/ERF transcription factors likely constitute a new susceptibility hub upregulated by TALEs and even non-TAL effectors upon infection with different pathogens.

Plants commonly harbour more than a hundred AP2/ERF transcription factors that generally have low sequence similarity and exert various functions, many of which are associated with resistance to various stress factors, which is especially true for the ERF group within the AP2/ERF family (Gutterson & Reuber, 2004; Licausi *et al*., 2013; Xie *et al*., 2019). Numerous studies have demonstrated that resistance to different pathogens, including several *Xanthomonas campestris* pathovars, is positively correlated with ERF expression levels in different plant species and increases upon ERF transgene overexpression (Gutterson & Reuber, 2004; Champion *et al*., 2009; Sherif *et al*., 2012; Licausi *et al*., 2013; Cacas *et al*., 2017; Tripathi *et al*., 2019). One of the type III effectors from *Xanthomonas euvesicatoria* has been shown to directly downregulate the ERF transcription factor in *Solanum lycopersicum* to promote susceptibility (Kim *et al*., 2013). At the molecular level, different ERFs induce the expression of multiple pathogenesis-related (PR) genes (Broekaert *et al*., 2006; Licausi *et al*., 2013). When this is taken into consideration, TALE-dependent activation of seemingly defence-associated ERF genes may seem counterintuitive.

The most obvious way for transcription factors to promote plant susceptibility is to downregulate the expression of defence-associated genes. Although AP2/ERFs function mainly as activators of downstream genes, some of them are transcriptional repressors that can attenuate the expression of defence-associated genes (Broekaert *et al*., 2006; Licausi *et al*., 2013; Xie *et al*., 2019). In addition to active transcriptional downregulation through specialized repressors, passive downregulation can also occur when a given transcription factor occupies a binding site to prevent interaction with a transcription activator (Licausi *et al*., 2013). Competition for binding sites has been shown for members of another large plant transcription factor family – the WRKY family (Turck *et al*., 2004). Despite their diversity, AP2/ERFs seem to interact with similar short GC-rich motifs (Licausi *et al*., 2013; Xie *et al*., 2019). Strong TALE-induced overproduction of a single ERF may lead to blocking of the corresponding cis-elements within the promoters of defence-associated genes that are normally regulated by other ERFs, thus disrupting the plant defence response.

Activation of plant defence pathways by ERFs can also be beneficial for Xanthomonas. It is now well-known that plant defence responses to biotic and abiotic stresses and to pathogens with different lifestyles often act antagonistically to each other (Kazan & Lyons, 2014; Li et al., 2019). Phytopathogenic bacteria often use different type III effectors to activate certain plant defence responses, which can lead to the repression of antagonistic defence pathways (Kazan & Lyons, 2014; Nakano & Mukaihara, 2019). Since ERFs are involved in various defence signalling pathways, the objective of TALE-dependent ERF activation may be the inhibition of defence responses most useful upon Xanthomonas attack through the misactivation of competing defence pathways. Indeed, overexpression of certain ERFs was shown to increase the susceptibility to one pathogen, with a simultaneous increase in resistance to another pathogen or even to an abiotic stress factor (Broekaert *et al*., 2006; Tsutsui et al., 2009; Li *et al*., 2018; Lu et al., 2020). As long as the molecular mechanism of action of most ERFs remains unknown, without further mechanistic studies, we can only speculate about possible defence pathways induced by TALE-activated ERFs. Generally, ERFs are the major mediators of ethylene signalling (Xie *et al*., 2019). Although ethylene plays an important role in plant defence against different pathogens, its role in plant-Xanthomonas interactions is controversial (van Loon *et al*., 2006; Shen *et al*., 2011; Kim *et al*., 2013). TALE-dependent activation of some ERFs can activate branches of the ethylene response that contribute more to susceptibility than to defence upon Xanthomonas attack.

Although in some cases it is known how TALE targets contribute to the plant susceptibility (Pérez-Quintero & Szurek, 2019), for the majority such explanation is still lacking. We believe that future high-throughput studies of effector-activated ERF regulons could reveal the molecular mechanisms underlying ERF-mediated plant susceptibility. It is likely that the clues can be found at the intersections of the regulons of different ERFs activated upon plant-Xanthomonas interactions.

## Acknowledgements

The authors acknowledge Aspen Orynbayev for providing Xcc strains. We apologize to our colleagues whose work was not cited in this article due to limited space. This study was supported by the RFBR [Grant No. 18-316-00134]; State task [Grant No. 0574-2019-0001]. The work was done using the scientific equipment of the Center for Collective Use «Biotechnology» at All-Russia Research Institute of Agricultural Biotechnology (Moscow, Russia; agreement RFMEFI62114×0003). This manuscript was edited for English language by Nature Research Editing Service.

## Author contributions

NZ and VT designed the research; NZ, ML, YM and VU performed the experiments; NZ and ML analyzed the data; NZ, ML and VT wrote the manuscript.

## References

Berardini TZ, Reiser L, Li D, Mezheritsky Y, Muller R, Strait E, Huala E. 2015. The Arabidopsis information resource: making and mining the “gold standard” annotated reference plant genome. Genesis 53: 474–485.

Bernard V, Brunaud V, Lecharny A. 2010. TC-motifs at the TATA-box expected position in plant genes: a novel class of motifs involved in the transcription regulation. BMC genomics 11: 166.

Broekaert WF, Delauré SL, De Bolle MF, Cammue BP. 2006. The role of ethylene in host-pathogen interactions. Annual Review of Phytopathology 44: 393–416.

Caarls L, Van der Does D, Hickman R, Jansen W, Verk MCV, Proietti S, Lorenzo O, Solano R, Pieterse CMJ, Van Wees SCM. 2017. Assessing the role of ETHYLENE RESPONSE FACTOR transcriptional repressors in salicylic acid-mediated suppression of jasmonic acid-responsive genes. Plant and Cell Physiology 58: 266–278.

Cacas JL, Pré M, Pizot M, Cissoko M, Diedhiou I, Jalloul A, Doumas P, Nicole M, Champion A. 2017. GhERF-IIb3 regulates the accumulation of jasmonate and leads to enhanced cotton resistance to blight disease. Molecular plant pathology 18: 825–836.

Champion A, Hebrard E, Parra B, Bournaud C, Marmey P, Tranchant C, Nicole M. 2009. Molecular diversity and gene expression of cotton ERF transcription factors reveal that group IXa members are responsive to jasmonate, ethylene and Xanthomonas. Molecular Plant Pathology 10: 471–485.

Cohn M, Morbitzer R, Lahaye T, Staskawicz BJ. 2016. Comparison of gene activation by two TAL effectors from X anthomonas axonopodis pv. manihotis reveals candidate host susceptibility genes in cassava. Molecular Plant Pathology 17: 875–889.

Denancé N, Szurek B, Doyle EL, Lauber E, Fontaine-Bodin L, Carrère S, Guy E, Hajri A, Cerutti A, Boureau T et al. 2018. Two ancestral genes shaped the Xanthomonas campestris TAL effector gene repertoire. New Phytologist 219: 391–407.

Erkes A, Mücke S, Reschke M, Boch J, Grau J. 2019. PrediTALE: A novel model learned from quantitative data allows for new perspectives on TALE targeting. PLoS computational biology 15: e1007206.

Erkes A, Reschke M, Boch J, Grau J. 2017. Evolution of transcription activator-like effectors in Xanthomonas oryzae. Genome biology and evolution 9: 1599–1615.

Feng JX, Liu D, Pan Y, Gong W, Ma LG, Luo JC, Deng XW, Zhu YX. 2005. An annotation update via cDNA sequence analysis and comprehensive profiling of developmental, hormonal or environmental responsiveness of the Arabidopsis AP2/EREBP transcription factor gene family. Plant molecular biology 59: 853–868.

Foucher J, Ruh M, Préveaux A, Carrère S, Pelletier S, Briand M, Serre RF, Jacques MA, Chen NW. 2020. Common bean resistance to Xanthomonas is associated with upregulation of the salicylic acid pathway and downregulation of photosynthesis. Research Square. doi: 10.21203/rs.3.rs-17010/v2

Goszczynska T, Serfontein JJ, Serfontein S. 2000. Introduction to Practical Phytobacteriology. ARC-Plant Protection Research Institute, Pretoria, South Africa.

Grau J, Reschke M, Erkes A, Streubel J, Morgan RD, Wilson GG, Koebnik R, Boch J. 2016. AnnoTALE: bioinformatics tools for identification, annotation, and nomenclature of TALEs from Xanthomonas genomic sequences. Scientific Reports 6: 21077.

Grau J, Wolf A, Reschke M, Bonas U, Posch S, Boch J. 2013. Computational predictions provide insights into the biology of TAL effector target sites. PLoS Comput Biol 9: e1002962.

Gutterson N, Reuber TL. 2004. Regulation of disease resistance pathways by AP2/ERF transcription factors. Current opinion in plant biology 7: 465–471.

Ha VTN, Dzhalilov FS, Vinogradova S, Kyrova E, Ignatov A. 2014. Genetic diversity of black rot pathogen in Russia: plant reaction. Zashchita kartofelya 2: 26–29. [Article in Russian]

Hommelsheim CM, Frantzeskakis L, Huang M, Ülker B. 2014. PCR amplification of repetitive DNA: a limitation to genome editing technologies and many other applications. Scientific reports 4: 1–13.

Hummel AW, Wilkins KE, Wang L, Cernadas RA, Bogdanove AJ. 2017. A transcription activator-like effector from Xanthomonas oryzae pv. oryzicola elicits dose-dependent resistance in rice. Molecular plant pathology 18: 55–66.

Hutin M, Pérez-Quintero AL, Lopez C, Szurek B. 2015. MorTAL Kombat: the story of defense against TAL effectors through loss-of-susceptibility. Frontiers in plant science 6: 535.

Kay S, Boch J, Bonas U. 2005. Characterization of AvrBs3-like effectors from a Brassicaceae pathogen reveals virulence and avirulence activities and a protein with a novel repeat architecture. Molecular plant-microbe interactions 18: 838–848.

Kazan K, Lyons R. 2014. Intervention of phytohormone pathways by pathogen effectors. The Plant Cell 26: 2285–2309.

Kim JG, Stork W, Mudgett MB. 2013. Xanthomonas type III effector XopD desumoylates tomato transcription factor SlERF4 to suppress ethylene responses and promote pathogen growth. Cell host & microbe 13: 143–154.

Li B, Ferreira MA, Huang M, Camargos LF, Yu X, Teixeira RM, Carpinetti PA, Mendes GC, Gouveia-Mageste BC, Liu C, et al. 2019. The receptor-like kinase NIK1 targets FLS2/BAK1 immune complex and inversely modulates antiviral and antibacterial immunity. Nature communications 10: 1–14.

Li Z, Tian Y, Xu J, Fu X, Gao J, Wang B, Peng R, Yao, Q. 2018. A tomato ERF transcription factor, SlERF84, confers enhanced tolerance to drought and salt stress but negatively regulates immunity against Pseudomonas syringae pv. tomato DC3000. Plant Physiology and Biochemistry 132: 683–695.

Licausi F, Ohme-Takagi M, Perata P. 2013. APETALA 2/Ethylene Responsive Factor (AP 2/ERF) transcription factors: Mediators of stress responses and developmental programs. New Phytologist 199: 639–649.

Lu W, Deng F, Jia J, Chen X, Li J, Wen Q, Li T, Meng Y, Shan W. 2020. The Arabidopsis thaliana gene AtERF019 negatively regulates plant resistance to Phytophthora parasitica by suppressing PAMP-triggered immunity. Molecular Plant Pathology 21: 1179–1193.

McGrath KC, Dombrecht B, Manners JM, Schenk PM, Edgar CI, Maclean DJ, Scheible W-R, Udvardi MK, Kazan K. 2005. Repressor-and activator-type ethylene response factors functioning in jasmonate signaling and disease resistance identified via a genome-wide screen of Arabidopsis transcription factor gene expression. Plant physiology 139: 949–959.

Mokryakov MV, Abdeev IA, Piruzyan ES, Schaad NW, Ignatov AN. 2010. Diversity of effector genes in plant pathogenic bacteria of genus Xanthomonas. Microbiology 79: 58–65.

Morbitzer R, Römer P, Boch J, Lahaye T. 2010. Regulation of selected genome loci using de novo-engineered transcription activator-like effector (TALE)-type transcription factors. Proceedings of the National Academy of Sciences 107: 21617–21622.

Mücke S, Reschke M, Erkes A, Schwietzer CA, Becker S, Streubel J, Morgan RD, Wilson GG, Grau J, Boch J. 2019. Transcriptional reprogramming of rice cells by Xanthomonas oryzae TALEs. Frontiers in plant science 10: 162.

Nakano M, Mukaihara T. 2019. Comprehensive Identification of PTI Suppressors in Type III Effector Repertoire Reveals that Ralstonia solanacearum Activates Jasmonate Signaling at Two Different Steps. International Journal of Molecular Sciences 20: 5992.

Okonechnikov K, Golosova O, Fursov M, Ugene Team. 2012. Unipro UGENE: a unified bioinformatics toolkit. Bioinformatics 28: 1166–1167.

Oliva R, Ji C, Atienza-Grande G, Huguet-Tapia JC, Pérez-Quintero A, Li T., Eom JS, Li C, Nguyen H, Liu B et al. 2019. Broad-spectrum resistance to bacterial blight in rice using genome editing. Nature biotechnology 37: 1344–1350.

Parkin IA, Koh C, Tang H, Robinson SJ, Kagale S, Clarke WE, Town CD, Nixon J, Krishnakumar V, Bidwell SL et al. 2014. Transcriptome and methylome profiling reveals relics of genome dominance in the mesopolyploid Brassica oleracea. Genome biology 15: 1–18.

Peng Z, Hu Y, Zhang J, Huguet-Tapia JC, Block AK, Park S, Sapkota S., Liu Z, Liu S, White FF. 2019. Xanthomonas translucens commandeers the host rate-limiting step in ABA biosynthesis for disease susceptibility. Proceedings of the National Academy of Sciences 116: 20938–20946.

Pereira AL, Carazzolle MF, Abe VY, de Oliveira ML, Domingues MN, Silva JC, Cernadas RA, Benedetti CE. 2014. Identification of putative TAL effector targets of the citrus canker pathogens shows functional convergence underlying disease development and defense response. BMC genomics 15: 1–15.

Pérez-Quintero AL, Lamy L, Gordon J, Escalon A, Cunnac S, Szurek B, Gagnevin L. 2015. QueTAL: a suite of tools to classify and compare TAL effectors functionally and phylogenetically. Frontiers in plant science 6: 545.

Pérez-Quintero AL, Rodriguez-R LM, Dereeper A, López C, Koebnik R, Szurek B, Cunnac S. 2013. An improved method for TAL effectors DNA-binding sites prediction reveals functional convergence in TAL repertoires of Xanthomonas oryzae strains. PloS one 8: e68464.

Pérez-Quintero AL, Szurek B. 2019. A decade decoded: spies and hackers in the history of TAL effectors research. Annual review of phytopathology 57: 459–481.

Pierce EJ, Rey MC. 2013. Assessing global transcriptome changes in response to South African cassava mosaic virus [ZA-99] infection in susceptible Arabidopsis thaliana. PloS one 8: e67534.

Postnikova OA, Minakova NY, Boutanaev AM, Nemchinov LG. 2011. Clustering of pathogen-response genes in the genome of Arabidopsis thaliana. Journal of integrative plant biology 53: 824–834.

Sayers EW, Beck J, Brister JR, Bolton EE, Canese K, Comeau DC, Funk K, Ketter A, Kim S, Kimchi A et al. 2020. Database resources of the national center for biotechnology information. Nucleic acids research 48: (D1), D9.

Schandry N, Jacobs JM, Szurek B, Pérez-Quintero AL. 2018. A cautionary TALE: how plant breeding may have favoured expanded TALE repertoires in Xanthomonas. Molecular plant pathology 19: 1297.

Shen X, Liu H, Yuan BIN, Li X, Xu C, Wang S. 2011. OsEDR1 negatively regulates rice bacterial resistance via activation of ethylene biosynthesis. Plant, cell & environment 34: 179–191.

Sherif S, El-Sharkawy I, Paliyath G, Jayasankar S. 2012. Differential expression of peach ERF transcriptional activators in response to signaling molecules and inoculation with Xanthomonas campestris pv. pruni. Journal of plant physiology 169: 731–739.

Streubel J, Baum H, Grau J, Stuttman J, Boch J. 2017. Dissection of TALE-dependent gene activation reveals that they induce transcription cooperatively and in both orientations. PloS one 12: e0173580.

Streubel J, Pesce C, Hutin M, Koebnik R, Boch J, Szurek B. 2013. Five phylogenetically close rice SWEET genes confer TAL effector-mediated susceptibility to Xanthomonas oryzae pv. oryzae. New Phytologist 200: 808–819.

Sugio A, Yang B, Zhu T, White FF. 2007. Two type III effector genes of Xanthomonas oryzae pv. oryzae control the induction of the host genes OsTFIIAγ1 and OsTFX1 during bacterial blight of rice. Proceedings of the National Academy of Sciences 104: 10720–10725.

Tariq R, Ji Z, Wang C, Tang Y, Zou L, Sun, H., Chen G, Zhao K. 2019. RNA-Seq analysis of gene expression changes triggered by Xanthomonas oryzae pv. oryzae in a susceptible rice genotype. Rice 12: 1–14.

Tran TT, Pérez-Quintero AL, Wonni I, Carpenter SC, Yu Y, Wang L, leach JE, Verdier V, Cunnac S, Bogdanove AJ et al. 2018. Functional analysis of African Xanthomonas oryzae pv. oryzae TALomes reveals a new susceptibility gene in bacterial leaf blight of rice. PLoS pathogens 14: e1007092.

Tripathi L, Tripathi JN, Shah T, Muiruri KS, Katari M. 2019. Molecular basis of disease resistance in banana progenitor Musa Balbisiana against Xanthomonas Campestris pv. Musacearum. Scientific reports 9: 1–17.

Truman W, de Zabala MT, Grant M. 2006. Type III effectors orchestrate a complex interplay between transcriptional networks to modify basal defence responses during pathogenesis and resistance. The Plant Journal 46: 14–33.

Tsutsui T, Kato W, Asada Y, Sako K, Sato T, Sonoda Y, Kidokoro S, Yamaguchi-Shinozaki K, Tamaoki M, Arakawa K et al. 2009. DEAR1, a transcriptional repressor of DREB protein that mediates plant defense and freezing stress responses in Arabidopsis. Journal of plant research 122: 633.

Turck F, Zhou A, Somssich IE. 2004. Stimulus-dependent, promoter-specific binding of transcription factor WRKY1 to its native promoter and the defense-related gene PcPR1-1 in parsley. The Plant Cell 16: 2573–2585.

van Loon LC, Geraats BP, Linthorst HJ. 2006. Ethylene as a modulator of disease resistance in plants. Trends in plant science 11: 184–191.

Verdier V, Triplett LR, Hummel AW, Corral R, Cernadas RA, Schmidt CL, Bogdanove AJ, Leach JE. 2012. Transcription activator-like (TAL) effectors targeting OsSWEET genes enhance virulence on diverse rice (Oryza sativa) varieties when expressed individually in a TAL effector□deficient strain of Xanthomonas oryzae. New Phytologist 196: 1197–1207.

Vicente JG, Holub EB. 2013. Xanthomonas campestris pv. campestris (cause of black rot of crucifers) in the genomic era is still a worldwide threat to brassica crops. Molecular plant pathology 14: 2–18.

Wang L, Rinaldi FC, Singh P, Doyle EL, Dubrow ZE, Tran TT, Pérez-Quintero AL, Szurek B, Bogdanove AJ. 2017. TAL effectors drive transcription bidirectionally in plants. Molecular plant 10: 285–296.

Wu D, von Roepenack-Lahaye E, Buntru M, de Lange O, Schandry N, Pérez-Quintero AL, Weinberg Z, Lowe-Power TM, Szurek B, Michael AJ et al. 2019. A plant pathogen type III effector protein subverts translational regulation to boost host polyamine levels. Cell Host & Microbe 26: 638–649.

Xie Z, Nolan TM, Jiang H, Yin Y. 2019. AP2/ERF transcription factor regulatory networks in hormone and abiotic stress responses in Arabidopsis. Frontiers in plant science 10: 228.

Zaka A, Grande G, Coronejo T, Quibod IL, Chen CW, Chang SJ, Szurek B, Arif M, Cruz CV, Oliva R. 2018. Natural variations in the promoter of OsSWEET13 and OsSWEET14 expand the range of resistance against Xanthomonas oryzae pv. oryzae. PloS one 13: e0203711.

