## Supplementary material for "An ERF transcription factor from Brassica oleracea: a new member of the emerging pathogenicity hub in plant-Xanthomonas interactions"

```
                           10        20        30        40        50        60        70
                  ....|....|....|....|....|....|....|....|....|....|....|....|....|....|
TalEN5_Xcc XY1-1  ATGGATCCCATTCGTTCGCGCACACCAAGTCCTGCCCGCGAGCTTCTGTCCGGACCCCAACCCGATGGGG
TalEN6_Xcc XY1-2  ATGGATCCCATTCGTTCGCGCACACCAAGTCCTGCCCGCGAGCTTCTGTCCGGACCCCAACCCGATGGGG
TalEN7_Xcc XY2-1  ATGGATCCCATTCGTTCGCGCACACCAAGTCCTGCCCGCGAGCTTCTGTCCGGACCCCAACCCGATGGGG
TalEN8_Xcc XY2-2  ATGGATCCCATTCGTTCGCGCACACCAAGTCCTGCCCGCGAGCTTCTGTCCGGACCCCAACCCGATGGGG
TalEN1_Xcc CN12   ATGGATCCCATTCGTTCGCGCACACCAAGTCCTGCCCGCGAGCTTCTGTCCGGACCCCAACCCGATGGGG
TalEN2_Xcc CN17   ATGGATCCCATTCGTTCGCGCACACCAAGTCCTGCCCGCGAGCTTCTGTCCGGACCCCAACCCGATGGGG
TalEN3_Xcc CN18   ATGGATCCCATTCGTTCGCGCACACCAAGTCCTGCCCGCGAGCTTCTGTCCGGACCCCAACCCGATGGGG

                           80        90       100       110       120       130       140
                  ....|....|....|....|....|....|....|....|....|....|....|....|....|....|
TalEN5_Xcc XY1-1  TTCAGCCGACTGCAGATCGTGGGGTGTCTCCGCCTGCCGGCGGCCCCCTGGATGGCTTGCCCGCTCGGCG
TalEN6_Xcc XY1-2  TTCAGCCGACTGCAGATCGTGGGGTGTCTCCGCCTGCCGGCGGCCCCCTGGATGGCTTGCCCGCTCGGCG
TalEN7_Xcc XY2-1  TTCAGCCGACTGCAGATCGTGGGGTGTCTCCGCCTGCCGGCGGCCCCCTGGATGGCTTGCCCGCTCGGCG
TalEN8_Xcc XY2-2  TTCAGCCGACTGCAGATCGTGGGGTGTCTCCGCCTGCCGGCGGCCCCCTGGATGGCTTGCCCGCTCGGCG
TalEN1_Xcc CN12   TTCAGCCGACTGCAGATCGTGGGGTGTCTCCGCCTGCCGGCGGCCCCCTGGATGGCTTGCCCGCTCGGCG
TalEN2_Xcc CN17   TTCAGCCGACTGCAGATCGTGGGGTGTCTCCGCCTGCCGGCGGCCCCCTGGATGGCTTGCCCGCTCGGCG
TalEN3_Xcc CN18   TTCAGCCGACTGCAGATCGTGGGGTGTCTCCGCCTGCCGGCGGCCCCCTGGATGGCTTGCCCGCTCGGCG

                          150       160       170       180       190       200       210
                  ....|....|....|....|....|....|....|....|....|....|....|....|....|....|
TalEN5_Xcc XY1-1  GACGATGTCCCGGACCCGGCTGCCATCTCCCCCTGCCCCCTCACCTGCGTTCTCGGCGGACAGCTTCAGT
TalEN6_Xcc XY1-2  GACGATGTCCCGGACCCGGCTGCCATCTCCCCCTGCCCCCTCACCTGCGTTCTCGGCGGACAGCTTCAGT
TalEN7_Xcc XY2-1  GACGATGTCCCGGACCCGGCTGCCATCTCCCCCTGCCCCCTCACCTGCGTTCTCGGCGGACAGCTTCAGT
TalEN8_Xcc XY2-2  GACGATGTCCCGGACCCGGCTGCCATCTCCCCCTGCCCCCTCACCTGCGTTCTCGGCGGACAGCTTCAGT
TalEN1_Xcc CN12   GACGATGTCCCGGACCCGGCTGCCATCTCCCCCTGCCCCCTCACCTGCGTTCTCGGCGGACAGCTTCAGT
TalEN2_Xcc CN17   GACGATGTCCCGGACCCGGCTGCCATCTCCCCCTGCCCCCTCACCTGCGTTCTCGGCGGACAGCTTCAGT
TalEN3_Xcc CN18   GACGATGTCCCGGACCCGGCTGCCATCTCCCCCTGCCCCCTCACCTGCGTTCTCGGCGGACAGCTTCAGT

                          220       230       240       250       260       270       280
                  ....|....|....|....|....|....|....|....|....|....|....|....|....|....|
TalEN5_Xcc XY1-1  GACCTGTTACGTCAGTTCGATCCGTCACTTTTTAATACATCGCTTTTTGATTCATTGCCTCCCTTCGGCG
TalEN6_Xcc XY1-2  GACCTGTTACGTCAGTTCGATCCGTCACTTTTTAATACATCGCTTTTTGATTCATTGCCTCCCTTCGGCG
TalEN7_Xcc XY2-1  GACCTGTTACGTCAGTTCGATCCGTCACTTTTTAATACATCGCTTTTTGATTCATTGCCTCCCTTCGGCG
TalEN8_Xcc XY2-2  GACCTGTTACGTCAGTTCGATCCGTCACTTTTTAATACATCGCTTTTTGATTCATTGCCTCCCTTCGGCG
TalEN1_Xcc CN12   GACCTGTTACGTCAGTTCGATCCGTCACTTTTTAATACATCGCTTTTTGATTCATTGCCTCCCTTCGGCG
TalEN2_Xcc CN17   GACCTGTTACGTCAGTTCGATCCGTCACTTTTTAATACATCGCTTTTTGATTCATTGCCTCCCTTCGGCG
TalEN3_Xcc CN18   GACCTGTTACGTCAGTTCGATCCGTCACTTTTTAATACATCGCTTTTTGATTCATTGCCTCCCTTCGGCG

                          290       300       310       320       330       340       350
                  ....|....|....|....|....|....|....|....|....|....|....|....|....|....|
TalEN5_Xcc XY1-1  CTCACCATACAGAGGCTGCCACAGGCGAGTGGGATGAGGTGCAATCGGGTCTGCGGGCAGCCGACGCCCC
TalEN6_Xcc XY1-2  CTCACCATACAGAGGCTGCCACAGGCGAGTGGGATGAGGTGCAATCGGGTCTGCGGGCAGCCGACGCCCC
TalEN7_Xcc XY2-1  CTCACCATACAGAGGCTGCCACAGGCGAGTGGGATGAGGTGCAATCGGGTCTGCGGGCAGCCGACGCCCC
TalEN8_Xcc XY2-2  CTCACCATACAGAGGCTGCCACAGGCGAGTGGGATGAGGTGCAATCGGGTCTGCGGGCAGCCGACGCCCC
TalEN1_Xcc CN12   CTCACCATACAGAGGCTGCCACAGGCGAGTGGGATGAGGTGCAATCGGGTCTGCGGGCAGCCGACGCCCC
TalEN2_Xcc CN17   CTCACCATACAGAGGCTGCCACAGGCGAGTGGGATGAGGTGCAATCGGGTCTGCGGGCAGCCGACGCCCC
TalEN3_Xcc CN18   CTCACCATACAGAGGCTGCCACAGGCGAGTGGGATGAGGTGCAATCGGGTCTGCGGGCAGCCGACGCCCC

                          360       370       380       390       400       410       420
                  ....|....|....|....|....|....|....|....|....|....|....|....|....|....|
TalEN5_Xcc XY1-1  CCCACCCACCATGCGCGTGGCTGTCACTGCCGCGCGGCCGCCGCGCGCCAAGCCGGCGCCGCGACGACGT
TalEN6_Xcc XY1-2  CCCACCCACCATGCGCGTGGCTGTCACTGCCGCGCGGCCGCCGCGCGCCAAGCCGGCGCCGCGACGACGT
TalEN7_Xcc XY2-1  CCCACCCACCATGCGCGTGGCTGTCACTGCCGCGCGGCCGCCGCGCGCCAAGCCGGCGCCGCGACGACGT
TalEN8_Xcc XY2-2  CCCACCCACCATGCGCGTGGCTGTCACTGCCGCGCGGCCGCCGCGCGCCAAGCCGGCGCCGCGACGACGT
TalEN1_Xcc CN12   CCCACCCACCATGCGCGTGGCTGTCACTGCCGCGCGGCCGCCGCGCGCCAAGCCGGCGCCGCGACGACGT
TalEN2_Xcc CN17   CCCACCCACCATGCGCGTGGCTGTCACTGCCGCGCGGCCGCCGCGCGCCAAGCCGGCGCCGCGACGACGT
TalEN3_Xcc CN18   CCCACCCACCATGCGCGTGGCTGTCACTGCCGCGCGGCCGCCGCGCGCCAAGCCGGCGCCGCGACGACGT

                          430       440       450       460       470       480       490
                  ....|....|....|....|....|....|....|....|....|....|....|....|....|....|
TalEN5_Xcc XY1-1  GCTGCGCAACCCTCCGACGCTTCGCCGGCCGCGCAGGTGGATCTACGCACGCTCGGCTACAGCCAGCAGC
TalEN6_Xcc XY1-2  GCTGCGCAACCCTCCGACGCTTCGCCGGCCGCGCAGGTGGATCTACGCACGCTCGGCTACAGCCAGCAGC
TalEN7_Xcc XY2-1  GCTGCGCAACCCTCCGACGCTTCGCCGGCCGCGCAGGTGGATCTACGCACGCTCGGCTACAGCCAGCAGC
TalEN8_Xcc XY2-2  GCTGCGCAACCCTCCGACGCTTCGCCGGCCGCGCAGGTGGATCTACGCACGCTCGGCTACAGCCAGCAGC
TalEN1_Xcc CN12   GCTGCGCAACCCTCCGACGCTTCGCCGGCCGCGCAGGTGGATCTACGCACGCTCGGCTACAGCCAGCAGC
TalEN2_Xcc CN17   GCTGCGCAACCCTCCGACGCTTCGCCGGCCGCGCAGGTGGATCTACGCACGCTCGGCTACAGCCAGCAGC
TalEN3_Xcc CN18   GCTGCGCAACCCTCCGACGCTTCGCCGGCCGCGCAGGTGGATCTACGCACGCTCGGCTACAGCCAGCAGC

                          500       510       520       530       540       550       560
                  ....|....|....|....|....|....|....|....|....|....|....|....|....|....|
TalEN5_Xcc XY1-1  AACAGGAGAAGATCAAACCGAAGGTTCGTTCGACAGTGGCGCAGCACCACGAGGCACTGGTCGGCCATGG
TalEN6_Xcc XY1-2  AACAGGAGAAGATCAAACCGAAGGTTCGTTCGACAGTGGCGCAGCACCACGAGGCACTGGTCGGCCATGG
TalEN7_Xcc XY2-1  AACAGGAGAAGATCAAACCGAAGGTTCGTTCGACAGTGGCGCAGCACCACGAGGCACTGGTCGGCCATGG
TalEN8_Xcc XY2-2  AACAGGAGAAGATCAAACCGAAGGTTCGTTCGACAGTGGCGCAGCACCACGAGGCACTGGTCGGCCATGG
TalEN1_Xcc CN12   AACAGGAGAAGATCAAACCGAAGGTTCGTTCGACAGTGGCGCAGCACCACGAGGCACTGGTCGGCCATGG
TalEN2_Xcc CN17   AACAGGAGAAGATCAAACCGAAGGTTCGTTCGACAGTGGCGCAGCACCACGAGGCACTGGTCGGCCATGG
TalEN3_Xcc CN18   AACAGGAGAAGATCAAACCGAAGGTTCGTTCGACAGTGGCGCAGCACCACGAGGCACTGGTCGGCCATGG

                          570       580       590       600       610       620       630
                  ....|....|....|....|....|....|....|....|....|....|....|....|....|....|
TalEN5_Xcc XY1-1  GTTTACACACGCGCACATCGTTGCGCTCAGCCAACACCCGGCAGCGTTAGGGACCGTCGCTGTCAAGTAT
TalEN6_Xcc XY1-2  GTTTACACACGCGCACATCGTTGCGCTCAGCCAACACCCGGCAGCGTTAGGGACCGTCGCTGTCAAGTAT
TalEN7_Xcc XY2-1  GTTTACACACGCGCACATCGTTGCGCTCAGCCAACACCCGGCAGCGTTAGGGACCGTCGCTGTCAAGTAT
TalEN8_Xcc XY2-2  GTTTACACACGCGCACATCGTTGCGCTCAGCCAACACCCGGCAGCGTTAGGGACCGTCGCTGTCAAGTAT
TalEN1_Xcc CN12   GTTTACACACGCGCACATCGTTGCGCTCAGCCAACACCCGGCAGCGTTAGGGACCGTCGCTGTCAAGTAT
TalEN2_Xcc CN17   GTTTACACACGCGCACATCGTTGCGCTCAGCCAACACCCGGCAGCGTTAGGGACCGTCGCTGTCAAGTAT
TalEN3_Xcc CN18   GTTTACACACGCGCACATCGTTGCGCTCAGCCAACACCCGGCAGCGTTAGGGACCGTCGCTGTCAAGTAT

                          640       650       660       670       680       690       700
                  ....|....|....|....|....|....|....|....|....|....|....|....|....|....|
TalEN5_Xcc XY1-1  CAGGACATGATCGCAGCGTTGCCAGAGGCGACACACGAAGCGATCGTTGGCGTCGGCAAACAGTGGTCCG
TalEN6_Xcc XY1-2  CAGGACATGATCGCAGCGTTGCCAGAGGCGACACACGAAGCGATCGTTGGCGTCGGCAAACAGTGGTCCG
TalEN7_Xcc XY2-1  CAGGACATGATCGCAGCGTTGCCAGAGGCGACACACGAAGCGATCGTTGGCGTCGGCAAACAGTGGTCCG
TalEN8_Xcc XY2-2  CAGGACATGATCGCAGCGTTGCCAGAGGCGACACACGAAGCGATCGTTGGCGTCGGCAAACAGTGGTCCG
TalEN1_Xcc CN12   CAGGACATGATCGCAGCGTTGCCAGAGGCGACACACGAAGCGATCGTTGGCGTCGGCAAACAGTGGTCCG
TalEN2_Xcc CN17   CAGGACATGATCGCAGCGTTGCCAGAGGCGACACACGAAGCGATCGTTGGCGTCGGCAAACAGTGGTCCG
TalEN3_Xcc CN18   CAGGACATGATCGCAGCGTTGCCAGAGGCGACACACGAAGCGATCGTTGGCGTCGGCAAACAGTGGTCCG

                          710       720       730       740       750       760       770
                  ....|....|....|....|....|....|....|....|....|....|....|....|....|....|
TalEN5_Xcc XY1-1  GCGCACGCGCCCTGGAGGCCTTGCTCACGGTGGCGGGAGAGTTGAGAGGTCCACCGTTACAGTTGGACAC
TalEN6_Xcc XY1-2  GCGCACGCGCCCTGGAGGCCTTGCTCACGGTGGCGGGAGAGTTGAGAGGTCCACCGTTACAGTTGGACAC
TalEN7_Xcc XY2-1  GCGCACGCGCCCTGGAGGCCTTGCTCACGGTGGCGGGAGAGTTGAGAGGTCCACCGTTACAGTTGGACAC
TalEN8_Xcc XY2-2  GCGCACGCGCCCTGGAGGCCTTGCTCACGGTGGCGGGAGAGTTGAGAGGTCCACCGTTACAGTTGGACAC
TalEN1_Xcc CN12   GCGCACGCGCCCTGGAGGCCTTGCTCACGGTGGCGGGAGAGTTGAGAGGTCCACCGTTACAGTTGGACAC
TalEN2_Xcc CN17   GCGCACGCGCCCTGGAGGCCTTGCTCACGGTGGCGGGAGAGTTGAGAGGTCCACCGTTACAGTTGGACAC
TalEN3_Xcc CN18   GCGCACGCGCTCTGGAGGCCTTGCTCACGGTGGCGGGAGAGTTGAGAGGTCCACCGTTACAGTTGGACAC

                          780       790       800       810       820       830       840
                  ....|....|....|....|....|....|....|....|....|....|....|....|....|....|
TalEN5_Xcc XY1-1  AGGCCAACTTCTCAAGATTGCAAAACGTGGCGGCGTGACCGCAGTGGAGGCAGTGCATGCATGGCGCAAT
TalEN6_Xcc XY1-2  AGGCCAACTTCTCAAGATTGCAAAACGTGGCGGCGTGACCGCAGTGGAGGCAGTGCATGCATGGCGCAAT
TalEN7_Xcc XY2-1  AGGCCAACTTCTCAAGATTGCAAAACGTGGCGGCGTGACCGCAGTGGAGGCAGTGCATGCATGGCGCAAT
TalEN8_Xcc XY2-2  AGGCCAACTTCTCAAGATTGCAAAACGTGGCGGCGTGACCGCAGTGGAGGCAGTGCATGCATGGCGCAAT
TalEN1_Xcc CN12   AGGCCAACTTCTCAAGATTGCAAAACGTGGCGGCGTGACCGCAGTGGAGGCAGTGCATGCATGGCGCAAT
TalEN2_Xcc CN17   AGGCCAACTTCTCAAGATTGCAAAACGTGGCGGCGTGACCGCAGTGGAGGCAGTGCATGCATGGCGCAAT
TalEN3_Xcc CN18   AGGCCAACTTCTCAAGATTGCAAAACGTGGCGGCGTGACCGCAGTGGAGGCAGTGCATGCATGGCGCAAT

                          850       860       870       880       890       900       910
                  ....|....|....|....|....|....|....|....|....|....|....|....|....|....|
TalEN5_Xcc XY1-1  GCACTGACGGGTGCCCCCCTGAACCTGACCCCGGAGCAGGTGGTGGCCATCGCCAGCAATATTGGTGGCA
TalEN6_Xcc XY1-2  GCACTGACGGGTGCCCCCCTGAACCTGACCCCGGAGCAGGTGGTGGCCATCGCCAGCAATATTGGTGGCA
TalEN7_Xcc XY2-1  GCACTGACGGGTGCCCCCCTGAACCTGACCCCGGAGCAGGTGGTGGCCATCGCCAGCAATATTGGTGGCA
TalEN8_Xcc XY2-2  GCACTGACGGGTGCCCCCCTGAACCTGACCCCGGAGCAGGTGGTGGCCATCGCCAGCAATATTGGTGGCA
TalEN1_Xcc CN12   GCACTGACGGGTGCCCCCCTGAACCTGACCCCGGAGCAGGTGGTGGCCATCGCCAGCAATATTGGTGGCA
TalEN2_Xcc CN17   GCACTGACGGGTGCCCCCCTGAACCTGACCCCGGAGCAGGTGGTGGCCATCGCCAGCAATATTGGTGGCA
TalEN3_Xcc CN18   GCACTGACGGGTGCCCCCCTGAACCTGACCCCGGAGCAGGTGGTGGCCATCGCCAGCAATATTGGTGGCA


                          920       930       940       950       960       970       980
                  ....|....|....|....|....|....|....|....|....|....|....|....|....|....|
TalEN5_Xcc XY1-1  AGCAGGCGCTGGAGACGGTGCAGGCGCTGTTGCCGGTGCTGTGCCAGGCCCATGGCCTGACCCCGCAGCA
TalEN6_Xcc XY1-2  AGCAGGCGCTGGAGACGGTGCAGGCGCTGTTGCCGGTGCTGTGCCAGGCCCATGGCCTGACCCCGCAGCA
TalEN7_Xcc XY2-1  AGCAGGCGCTGGAGACGGTGCAGGCGCTGTTGCCGGTGCTGTGCCAGGCCCATGGCCTGACCCCGCAGCA
TalEN8_Xcc XY2-2  AGCAGGCGCTGGAGACGGTGCAGGCGCTGTTGCCGGTGCTGTGCCAGGCCCATGGCCTGACCCCGCAGCA
TalEN1_Xcc CN12   AGCAGGCGCTGGAGACGGTGCAGGCGCTGTTGCCGGTGCTGTGCCAGGCCCATGGCCTGACCCCGGAGCA
TalEN2_Xcc CN17   AGCAGGCGCTGGAGACGGTGCAGGCGCTGTTGCCGGTGCTGTGCCAGGCCCATGGCCTGACCCCGGAGCA
TalEN3_Xcc CN18   AGCAGGCGCTGGAGACGGTGCAGGCGCTGTTGCCGGTGCTGTGCCAGGCCCATGGCCTGACCCCGCAGCA

                          990       1000      1010      1020      1030      1040      1050
                  ....|....|....|....|....|....|....|....|....|....|....|....|....|....|
TalEN5_Xcc XY1-1  GGTGGTGGCCATCGCCAGCAATGGCGGTGGCAAGCAGGCGCTGGAGACGGTGCAGCGGCTGTTGCCGGTG
TalEN6_Xcc XY1-2  GGTGGTGGCCATCGCCAGCAATGGCGGTGGCAAGCAGGCGCTGGAGACGGTGCAGCGGCTGTTGCCGGTG
TalEN7_Xcc XY2-1  GGTGGTGGCCATCGCCAGCAATGGCGGTGGCAAGCAGGCGCTGGAGACGGTGCAGCGGCTGTTGCCGGTG
TalEN8_Xcc XY2-2  GGTGGTGGCCATCGCCAGCAATGGCGGTGGCAAGCAGGCGCTGGAGACGGTGCAGCGGCTGTTGCCGGTG
TalEN1_Xcc CN12   GGTGGTGGCCATCGCCAGCAATGGCGGTGGCAAGCAGGCGCTGGAGACGGTGCAGCGGCTGTTGCCGGTG
TalEN2_Xcc CN17   GGTGGTGGCCATCGCCAGCAATGGCGGTGGCAAGCAGGCGCTGGAGACGGTGCAGCGGCTGTTGCCGGTG
TalEN3_Xcc CN18   GGTGGTGGCCATCGCCAGCCACGATGGCGGCAAGCAGGCGCTGGAGACGGTGCAGCGGCTGTTGCCGGTG

                          1060      1070      1080      1090      1100      1110      1120
                  ....|....|....|....|....|....|....|....|....|....|....|....|....|....|
TalEN5_Xcc XY1-1  CTGTGCCAGGCCCATGGCCTGACCCCGGAGCAGGTGGTGGCCATCGCCAGCAATATTGGTGGCAAGCAGG
TalEN6_Xcc XY1-2  CTGTGCCAGGCCCATGGCCTGACCCCGGAGCAGGTGGTGGCCATCGCCAGCAATATTGGTGGCAAGCAGG
TalEN7_Xcc XY2-1  CTGTGCCAGGCCCATGGCCTGACCCCGGAGCAGGTGGTGGCCATCGCCAGCAATATTGGTGGCAAGCAGG
TalEN8_Xcc XY2-2  CTGTGCCAGGCCCATGGCCTGACCCCGGAGCAGGTGGTGGCCATCGCCAGCAATATTGGTGGCAAGCAGG
TalEN1_Xcc CN12   CTGTGCCAGGCCCATGGCCTGACCCCGGAGCAGGTGGTGGCCATCGCCAGCAATATTGGTGGCAAGCAGG
TalEN2_Xcc CN17   CTGTGCCAGGCCCATGGCCTGACCCCGGAGCAGGTGGTGGCCATCGCCAGCAATATTGGTGGCAAGCAGG
TalEN3_Xcc CN18   CTGTGCCAGGCCCATGGCCTGACCCCGCAGCAGGTGGTGGCCATCGCCAGCCACGATGGCGGCAAGCAGG

                          1130      1140      1150      1160      1170      1180      1190
                  ....|....|....|....|....|....|....|....|....|....|....|....|....|....|
TalEN5_Xcc XY1-1  CGCTGGAGACGGTGCAGGCGCTGTTGCCGGTGCTGTGCCAGGCCCATGGCCTGACCCCGCAGCAGGTGGT
TalEN6_Xcc XY1-2  CGCTGGAGACGGTGCAGGCGCTGTTGCCGGTGCTGTGCCAGGCCCATGGCCTGACCCCGCAGCAGGTGGT
TalEN7_Xcc XY2-1  CGCTGGAGACGGTGCAGGCGTTGTTGCCGGTGCTGTGCCAGGCCCATGGCCTGACCCCGCAGCAGGTGGT
TalEN8_Xcc XY2-2  CGCTGGAGACGGTGCAGGCGCTGTTGCCGGTGCTGTGCCAGGCCCATGGCCTGACCCCGCAGCAGGTGGT
TalEN1_Xcc CN12   CGCTGGAGACGGTGCAGGCGCTGTTGCCGGTGCTGTGCCAGGCCCATGGCCTGACCCCGCAGCAGGTGGT
TalEN2_Xcc CN17   CGCTGGAGACGGTGCAGGCGCTGTTGCCGGTGCTGTGCCAGGCCCATGGCCTGACCCCGCAGCAGGTGGT
TalEN3_Xcc CN18   CGCTGGAGACGGTGCAGCGGCTGTTGCCGGTGCTGTGCCAGGCCCATGGCCTGACCCCGCAGCAGGTGGT

                          1200      1210      1220      1230      1240      1250      1260
                  ....|....|....|....|....|....|....|....|....|....|....|....|....|....|
TalEN5_Xcc XY1-1  GGCCATCGCCAGCAATGGCGGTGGCAAGCAGGCGCTGGAGACGGTGCAGCGGCTGTTGCCGGTGCTGTGC
TalEN6_Xcc XY1-2  GGCCATCGCCAGCAATGGCGGTGGCAAGCAGGCGCTGGAGACGGTGCAGCGGCTGTTGCCGGTGCTGTGC
TalEN7_Xcc XY2-1  GGCCATCGCCAGCAATGGCGGTGGCAAGCAGGCGCTGGAGACGGTGCAGCGGCTGTTGCCGGTGCTGTGC
TalEN8_Xcc XY2-2  GGCCATCGCCAGCAATGGCGGTGGCAAGCAGGCGCTGGAGACGGTGCAGCGGCTGTTGCCGGTGCTGTGC
TalEN1_Xcc CN12   GGCCATCGCCAGCAATGGCGGTGGCAAGCAGGCGCTGGAGACGGTGCAGCGGCTGTTGCCGGTGCTGTGC
TalEN2_Xcc CN17   GGCCATCGCCAGCAATGGCGGTGGCAAGCAGGCGCTGGAGACGGTGCAGCGGCTGTTGCCGGTGCTGTGC
TalEN3_Xcc CN18   GGCCATCGCCAGCAATGGCGGTGGCAAGCAGGCGCTGGAGACGGTGCAGCGGCTGTTGCCGGTGCTGTGC

                          1270      1280      1290      1300      1310      1320      1330
                  ....|....|....|....|....|....|....|....|....|....|....|....|....|....|
TalEN5_Xcc XY1-1  CAGGCCCATGGCCTGACCCCGGAGCAGGTGGTGGCCATCGCCAGCAATATTGGTGGCAAGCAGGCGCTGG
TalEN6_Xcc XY1-2  CAGGCCCATGGCCTGACCCCGGAGCAGGTGGTGGCCATCGCCAGCAATATTGGTGGCAAGCAGGCGCTGG
TalEN7_Xcc XY2-1  CAGGCCCATGGCCTGACCCCGGAGCAGGTGGTGGCCATCGCCAGCAATATTGGTGGCAAGCAGGCGCTGG
TalEN8_Xcc XY2-2  CAGGCCCATGGCCTGACCCCGGAGCAGGTGGTGGCCATCGCCAGCAATATTGGTGGCAAGCAGGCGCTGG
TalEN1_Xcc CN12   CAGGCCCATGGCCTGACCCCGGAGCAGGTGGTGGCCATCGCCAGCAATATTGGTGGCAAGCAGGCGCTGG
TalEN2_Xcc CN17   CAGGCCCATGGCCTGACCCCGGAGCAGGTGGTGGCCATCGCCAGCAATATTGGTGGCAAGCAGGCGCTGG
TalEN3_Xcc CN18   CAGGCCCATGGCCTGACCCCGGAGCAGGTGGTGGCCATCGCCAGCAATAGCGGTGGCAAGCAGGCGCTGG

                          1340      1350      1360      1370      1380      1390      1400
                  ....|....|....|....|....|....|....|....|....|....|....|....|....|....|
TalEN5_Xcc XY1-1  AGACGGTGCAGCGGCTGTTGCCGGTGCTGTGCCAGGCCCCCCATGACCTGACCCCGGAGCAGGTGGTGGC
TalEN6_Xcc XY1-2  AGACGGTGCAGCGGCTGTTGCCGGTGCTGTGCCAGGCCCCCCATGACCTGACCCCGGAGCAGGTGGTGGC
TalEN7_Xcc XY2-1  AGACGGTGCAGCGGCTGTTGCCGGTGCTGTGCCAGGCCCCCCATGACCTGACCCCGGAGCAGGTGGTGGC
TalEN8_Xcc XY2-2  AGACGGTGCAGCGGCTGTTGCCGGTGCTGTGCCAGGCCCCCCATGACCTGACCCCGGAGCAGGTGGTGGC
TalEN1_Xcc CN12   AGACGGTGCAGGCGCTGTTGCCGGTGCTGTGCCAGGCCC---ATGGCCTGACCCCGGAGCAGGTGGTGGC
TalEN2_Xcc CN17   AGACGGTGCAGGCGCTGTTGCCGGTGCTGTGCCAGGCCC---ATGGCCTGACCCCGGAGCAGGTGGTGGC
TalEN3_Xcc CN18   AGACGGTGCAGGCGCTGTTGCCGGTGCTGTGCCAGGCCC---ATGGCCTGACCCCGCAGCAGGTGGTGGC
                          1410      1420      1430      1440      1450      1460      1470
                  ....|....|....|....|....|....|....|....|....|....|....|....|....|....|
TalEN5_Xcc XY1-1  CATCGCCAGCAATATTGGTGGCAAGCAGGCGCTGGAGACGGTGCAGGCGCTGTTGCCGGTGCTGTGCCAG
TalEN6_Xcc XY1-2  CATCGCCAGCAATATTGGTGGCAAGCAGGCGCTGGAGACGGTGCAGGCGCTGTTGCCGGTGCTGTGCCAG
TalEN7_Xcc XY2-1  CATCGCCAGCAATATTGGTGGCAAGCAGGCGCTGGAGACGGTGCAGGCGCTGTTGCCGGTGCTGTGCCAG
TalEN8_Xcc XY2-2  CATCGCCAGCAATATTGGTGGCAAGCAGGCGCTGGAGACGGTGCAGGCGCTGTTGCCGGTGCTGTGCCAG
TalEN1_Xcc CN12   CATCGCCAGCAATATTGGTGGCAAGCAGGCGCTGGAGACGGTGCAGGCGCTGTTGCCGGTGCTGTGCCAG
TalEN2_Xcc CN17   CATCGCCAGCAATATTGGTGGCAAGCAGGCGCTGGAGACGGTGCAGGCGCTGTTGCCGGTGCTGTGCCAG
TalEN3_Xcc CN18   CATCGCCAGCAATAGCGGTGGCAAGCAGGCGCTGGAGACGGTGCAGGCGCTGTTGCCGGTGCTGTGCCAG

                          1480      1490      1500      1510      1520      1530      1540
                  ....|....|....|....|....|....|....|....|....|....|....|....|....|....|
TalEN5_Xcc XY1-1  GCCCATGGCCTGACCCCGGAGCAGGTGGTGGCCATCGCCAGCAATATTGGTGGCAAGCAGGCGCTGGAGA
TalEN6_Xcc XY1-2  GCCCATGGCCTGACCCCGGAGCAGGTGGTGGCCATCGCCAGCAATATTGGTGGCAAGCAGGCGCTGGAGA
TalEN7_Xcc XY2-1  GCCCATGGCCTGACCCCGGAGCAGGTGGTGGCCATCGCCAGCAATATTGGTGGCAAGCAGGCGCTGGAGA
TalEN8_Xcc XY2-2  GCCCATGGCCTGACCCCGGAGCAGGTGGTGGCCATCGCCAGCAATATTGGTGGCAAGCAGGCGCTGGAGA
TalEN1_Xcc CN12   GCCCATGGCCTGACCCCGGAGCAGGTGGTGGCCATCGCCAGCAATATTGGTGGCAAGCAGGCGCTGGAGA
TalEN2_Xcc CN17   GCCCATGGCCTGACCCCGGAGCAGGTGGTGGCCATCGCCAGCAATATTGGTGGCAAGCAGGCGCTGGAGA
TalEN3_Xcc CN18   GCCCATGGCCTGACCCCGGAGCAGGTGGTGGCCATCGCCAGCAATATTGGTGGCAAGCAGGCGCTGGAGA

                          1550      1560      1570      1580      1590      1600      1610
                  ....|....|....|....|....|....|....|....|....|....|....|....|....|....|
TalEN5_Xcc XY1-1  CGGTGCAGCGGCTGTTGCCGGTGCTGTGCCAGGCCCATGGCCTGACCCCGGAGCAGGTGGTGGCCATCGC
TalEN6_Xcc XY1-2  CGGTGCAGCGGCTGTTGCCGGTGCTGTGCCAGGCCCATGGCCTGACCCCGGAGCAGGTGGTGGCCATCGC
TalEN7_Xcc XY2-1  CGGTGCAGCGGCTGTTGCCGGTGCTGTGCCAGGCCCATGGCCTGACCCCGGAGCAGGTGGTGGCCATCGC
TalEN8_Xcc XY2-2  CGGTGCAGCGGCTGTTGCCGGTGCTGTGCCAGGCCCATGGCCTGACCCCGGAGCAGGTGGTGGCCATCGC
TalEN1_Xcc CN12   CGGTGCAGCGGCTGTTGCCGGTGCTGTGCCAGGCCCATGGCCTGACCCCGGAGCAGGTGGTGGCCATCGC
TalEN2_Xcc CN17   CGGTGCAGCGGCTGTTGCCGGTGCTGTGCCAGGCCCATGGCCTGACCCCGGAGCAGGTGGTGGCCATCGC
TalEN3_Xcc CN18   CGGTGCAGCGGCTGTTGCCGGTGCTGTGCCAGGCCCATGGCCTGACCCCGGAGCAGGTGGTGGCCATCGC

                          1620      1630      1640      1650      1660      1670      1680
                  ....|....|....|....|....|....|....|....|....|....|....|....|....|....|
TalEN5_Xcc XY1-1  CAGCCATGATGGCGGCAAGCAGGCGCTGGAGACGGTGCAGCGGCTGTTGCCGGTGCTGTGCCAGGCCCAT
TalEN6_Xcc XY1-2  CAGCCACGATGGCGGCAAGCAGGCGCTGGAGACGGTGCAGCGGCTGTTGCCGGTGCTGTGCCAGGCCCAT
TalEN7_Xcc XY2-1  CAGCCACCACGGCGGCAAGCAGGCGCTGGAGACGGTGCAGCGGCTGTTGCCGGTGCTGTGCCAGGCCCAT
TalEN8_Xcc XY2-2  CAGCCACGATGGCGGCAAGCAGGCGCTGGAGACGGTGCAGCGGCTGTTGCCGGTGCTGTGCCAGGCCCAT
TalEN1_Xcc CN12   CAGCCACGATGGCGGCAAGCAGGCGCTGGAGACGGTGCAGCGGCTGTTGCCGGTGCTGTGCCAGGCCCAT
TalEN2_Xcc CN17   CAGCCACGATGGCGGCAAGCAGGCGCTGGAGACGGTGCAGCGGCTGTTGCCGGTGCTGTGCCAGGCCCAT
TalEN3_Xcc CN18   CAGCCACGATGGCGGCAAGCAGGCGCTGGAGACGGTGCAGCGGCTGTTGCCGGTGCTGTGCCAGGCCCAT

                          1690      1700      1710      1720      1730      1740      1750
                  ....|....|....|....|....|....|....|....|....|....|....|....|....|....|
TalEN5_Xcc XY1-1  GGCCTGACCCCGGAGCAGGTGGTGGCCATCGCCAGCCACGATGGCGGCAAGCAGGCGCTGGAGACGGTGC
TalEN6_Xcc XY1-2  GGCCTGACCCCGGAGCAGGTGGTGGCCATCGCCAGCCACGATGGCGGCAAGCAGGCGCTGGAGACGGTGC
TalEN7_Xcc XY2-1  GGCCTGACCCCGGAGCAGGTGGTGGCCATCGCCAGCCACGATGGCGGCAAGCAGGCGCTGGAGACGGTGC
TalEN8_Xcc XY2-2  GGCCTGACCCCGGAGCAGGTGGTGGCCATCGCCAGCCACGATGGCGGCAAGCAGGCGCTGGAGACGGTGC
TalEN1_Xcc CN12   GGCCTGACCCCGGAGCAGGTGGTGGCCATCGCCAGCCACGATGGCGGCAAGCAGGCGCTGGAGACGGTGC
TalEN2_Xcc CN17   GGCCTGACCCCGGAGCAGGTGGTGGCCATCGCCAGCCACGATGGCGGCAAGCAGGCGCTGGAGACGGTGC
TalEN3_Xcc CN18   GGCCTGACCCCGCAGCAGGTGGTGGCCATCGCCAGCAATGGCGGTGGCAAGCAGGCGCTGGAGACGGTGC

                          1760      1770      1780      1790      1800      1810      1820
                  ....|....|....|....|....|....|....|....|....|....|....|....|....|....|
TalEN5_Xcc XY1-1  AGCGGCTGTTGCCGGTGCTGTGCCAGGCCCATGGCCTGACCCCGGAGCAGGTGGTGGCCATCGCCAGCCA
TalEN6_Xcc XY1-2  AGCGGCTGTTGCCGGTGCTGTGCCAGGCCCATGGCCTGACCCCGGAGCAGGTGGTGGCCATCGCCAGCCA
TalEN7_Xcc XY2-1  AGCGGCTGTTGCCGGTGCTGTGCCAGGCCCATGGCCTGACCCCGGAGCAGGTGGTGGCCATCGCCAGCCA
TalEN8_Xcc XY2-2  AGCGGCTGTTGCCGGTGCTGTGCCAGGCCCATGGCCTGACCCCGGAGCAGGTGGTGGCCATCGCCAGCCA
TalEN1_Xcc CN12   AGCGGCTGTTGCCGGTGCTGTGCCAGGCCCATGACCTGACCCCGGAGCAGGTGGTGGCCATCGCCAGCCA
TalEN2_Xcc CN17   AGCGGCTGTTGCCGGTGCTGTGCCAGGCCCATGACCTGACCCCGGAGCAGGTGGTGGCCATCGCCAGCCA
TalEN3_Xcc CN18   AGCGGCTGTTGCCGGTGCTGTGCCAGGCCCATGGCCTGACCCCGCAGCAGGTGGTGGCCATCGCCAGCAA

                          1830      1840      1850      1860      1870      1880      1890
                  ....|....|....|....|....|....|....|....|....|....|....|....|....|....|
TalEN5_Xcc XY1-1  CGATGGCGGCAAGCAGGCGCTGGAGACGGTGCAGCGGCTGTTGCCGGTGCTGTGCCAGGCCCATGGCCTG
TalEN6_Xcc XY1-2  CGATGGCGGCAAGCAGGCGCTGGAGACGGTGCAGCGGCTGTTGCCGGTGCTGTGCCAGGCCCATGGCCTG
TalEN7_Xcc XY2-1  CGATGGCGGCAAGCAGGCGCTGGAGACGGTGCAGCGGCTGTTGCCGGTGCTGTGCCAGGCCCATGGCCTG
TalEN8_Xcc XY2-2  CGATGGCGGCAAGCAGGCGCTGGAGACGGTGCAGCGGCTGTTGCCGGTGCTGTGCCAGGCCCATGGCCTG
TalEN1_Xcc CN12   CGATGGCGGCAAGCAGGCGCTGGAGACGGTGCAGCGGCTGTTGCCGGTGCTGTGCCAGGCCCATGGCCTG
TalEN2_Xcc CN17   CGATGGCGGCAAGCAGGCGCTGGAGACGGTGCAGCGGCTGTTGCCGGTGCTGTGCCAGGCCCATGGCCTG
TalEN3_Xcc CN18   TAGCGGTGGCAAGCAGGCGCTGGAGACGGTGCAGGCGCTGTTGCCGGTGCTGTGCCAGGCCCATGGCCTG

                          1900      1910      1920      1930      1940      1950      1960
                  ....|....|....|....|....|....|....|....|....|....|....|....|....|....|
TalEN5_Xcc XY1-1  ACCCCGGAGCAGGTGGTGGCCATCGCCAGCAATGGCGGTGGCAAGCAGGCGCTGGAGACGGTGCAGCGGC
TalEN6_Xcc XY1-2  ACCCCGGAGCAGGTGGTGGCCATCGCCAGCAATGGCGGTGGCAAGCAGGCGCTGGAGACGGTGCAGCGGC
TalEN7_Xcc XY2-1  ACCCCGGAGCAGGTGGTGGCCATCGCCAGCAATGGCGGTGGCAAGCAGGCGCTGGAGACGGTGCAGCGGC
TalEN8_Xcc XY2-2  ACCCCGGAGCAGGTGGTGGCCATCGCCAGCAATGGCGGTGGCAAGCAGGCGCTGGAGACGGTGCAGCGGC
TalEN1_Xcc CN12   ACCCCGGAGCAGGTGGTGGCCATCGCCAGCAATGGCGGTGGCAAGCAGGCGCTGGAGACGGTGCAGCGGC
TalEN2_Xcc CN17   ACCCCGGAGCAGGTGGTGGCCATCGCCAGCAATGGCGGTGGCAAGCAGGCGCTGGAGACGGTGCAGCGGC
TalEN3_Xcc CN18   ACCCCGCAGCAGGTGGTGGCCATCGCCAGCAATAGCGGTGGCAAGCAGGCGCTGGAGACGGTGCAGGCGC

                          1970      1980      1990      2000      2010      2020      2030
                  ....|....|....|....|....|....|....|....|....|....|....|....|....|....|
TalEN5_Xcc XY1-1  TGTTGCCGGTGCTGTGCCAGGCCCATGGCCTGACCCCGGAGCAGGTGGTGGCCATCGCCAGCAATATTGG
TalEN6_Xcc XY1-2  TGTTGCCGGTGCTGTGCCAGGCCCATGGCCTGACCCCGGAGCAGGTGGTGGCCATCGCCAGCAATATTGG
TalEN7_Xcc XY2-1  TGTTGCCGGTGCTGTGCCAGGCCCATGGCCTGACCCCGGAGCAGGTGGTGGCCATCGCCAGCAATATTGG
TalEN8_Xcc XY2-2  TGTTGCCGGTGCTGTGCCAGGCCCATGGCCTGACCCCGGAGCAGGTGGTGGCCATCGCCAGCAATATTGG
TalEN1_Xcc CN12   TGTTGCCGGTGCTGTGCCAGGCCCATGGCCTGACCCCGGAGCAGGTGGTGGCCATCGCCAGCAATATTGG
TalEN2_Xcc CN17   TGTTGCCGGTGCTGTGCCAGGCCCATGGCCTGACCCCGGAGCAGGTGGTGGCCATCGCCAGCAATATTGG
TalEN3_Xcc CN18   TGTTGCCGGTGCTGTGCCAGGCCCATGGCCTGACCCCGGAGCAGGTGGTGGCCATCGCCAGCAATATTGG

                          2040      2050      2060      2070      2080      2090      2100
                  ....|....|....|....|....|....|....|....|....|....|....|....|....|....|
TalEN5_Xcc XY1-1  TGGCAAGCAGGCGCTGGAGACGGTGCAGGCGCTGTTGCCGGTGCTGTGCCAGGCCCATGGCCTGACCCCG
TalEN6_Xcc XY1-2  TGGCAAGCAGGCGCTGGAGACGGTGCAGCGGCTGTTGCCGGTGCTGTGCCAGGCCCATGGCCTGACCCCG
TalEN7_Xcc XY2-1  TGGCAAGCAGGCGCTGGAGACGGTGCAGGCGCTGTTGCCGGTGCTGTGCCAGGCCCATGGCCTGACCCCG
TalEN8_Xcc XY2-2  TGGCAAGCAGGTGCTGGAGACGGTGCAGGCGCTGTTGCCGGTGCTGTGCCAGGCCCATGGCCTGACCCCG
TalEN1_Xcc CN12   TGGCAAGCAGGCGCTGGAGACGGTGCAGGCGCTGTTGCCGGTGCTGTGCCAGGCCCATGGCCTGACCCCG
TalEN2_Xcc CN17   TGGCAAGCAGGCGCTGGAGACGGTGCAGGCGCTGTTGCCGGTGCTGTGCCAGGCCCATGGCCTGACCCCG
TalEN3_Xcc CN18   TGGCAAGCAGGCGCTGGAGACGGTGCAGCGGCTGTTGCCGGTGCTGTGCCAGGCCCATGGCCTGACCCCG


                          2110      2120      2130      2140      2150      2160      2170
                  ....|....|....|....|....|....|....|....|....|....|....|....|....|....|
TalEN5_Xcc XY1-1  GAGCAGGTGGTGGCCATCGCCAGCAATATTGGTGGCAAGCAGGCGCTGGAGACGGTGCAGGCGCTGTTGC
TalEN6_Xcc XY1-2  GAGCAGGTGGTGGCCATCGCCAGCAATATTGGTGGCAAGCAGGCGCTGGAGACGGTGCAGGCGCTGTTGC
TalEN7_Xcc XY2-1  GAGCAGGTGGTGGCCATCGCCAGCAATATTGGTGGCAAGCAGGCGCTGGAGACGGTGCAGGCGCTGTTGC
TalEN8_Xcc XY2-2  GAGCAGGTGGTGGCCATCGCCAGCAATATTGGTGGCAAGCAGGCGCTGGAGACGGTGCAGGCGCTGTTGC
TalEN1_Xcc CN12   GAGCAGGTGGTGGCCATCGCCAGCAATATTGGTGGCAAGCAGGCGCTGGAGACGGTGCAGGCGCTGTTGC
TalEN2_Xcc CN17   GAGCAGGTGGTGGCCATCGCCAGCAATATTGGTGGCAAGCAGGCGCTGGAGACGGTGCAGGCGCTGTTGC
TalEN3_Xcc CN18   GAGCAGGTGGTGGCCATCGCCAGCAATATTGGTGGCAAGCAGGCGCTGGAGACGGTGCAGCGGCTGTTGC

                          2180      2190      2200      2210      2220      2230      2240
                  ....|....|....|....|....|....|....|....|....|....|....|....|....|....|
TalEN5_Xcc XY1-1  CGGTGCTGTGCCAGGCCCATGGCCTGACCCCGGAGCAGGTGGTGGCCATCGCCAGCAATATTGGTGGCAA
TalEN6_Xcc XY1-2  CGGTGCTGTGCCAGGCCCATGGCCTGACCCCGGAGCAGGTGGTGGCCATCGCCAGCAATATTGGTGGCAA
TalEN7_Xcc XY2-1  CGGTGCTGTGCCAGGCCCATGGCCTGACCCCGGAGCAGGTGGTGGCCATCGCCAGCAATATTGGTGGCAA
TalEN8_Xcc XY2-2  CGGTGCTGTGCCAGGCCCATGGCCTGACCCCGGAGCAGGTGGTGGCCATCGCCAGCAATATTGGTGGCAA
TalEN1_Xcc CN12   CGGTGCTGTGCCAGGCCCATGGCCTGACCCCGGAGCAGGTGGTGGCCATCGCCAGCAATATTGGTGGCAA
TalEN2_Xcc CN17   CGGTGCTGTGCCAGGCCCATGGCCTGACCCCGGAGCAGGTGGTGGCCATCGCCAGCAATATTGGTGGCAA
TalEN3_Xcc CN18   CGGTGCTGTGCCAGGCCCATGGCCTGACCCCGGAGCAGGTGGTGGCCATCGCCAGCAATATTGGTGGCAA

                          2250      2260      2270      2280      2290      2300      2310
                  ....|....|....|....|....|....|....|....|....|....|....|....|....|....|
TalEN5_Xcc XY1-1  GCAGGCGCTGGAGACGGTGCAGGCGCTGTTGCCGGTGCTGTGCCAGGCCCATGGCCTGACCCCGGAGCAG
TalEN6_Xcc XY1-2  GCAGGCGCTGGAGACGGTGCAGGCGCTGTTGCCGGTGCTGTGCCAGGCCCATGGCCTGACCCCGGAGCAG
TalEN7_Xcc XY2-1  GCAGGCGCTGGAGACGGTGCAGGCGCTGTTGCCGGTGCTGTGCCAGGCCCATGGCCTGACCCCGGAGCAG
TalEN8_Xcc XY2-2  GCAGGCGCTGGAGACGGTGCAGGCGCTGTTGCCGGTGCTGTGCCAGGCCCATGGCCTGACCCCGGAGCAG
TalEN1_Xcc CN12   GCAGGCGCTGGAGACGGTGCAGCGGCTGTTGCCGGTGCTGTGCCAGGCCCATGGCCTGACCCCGGAGCAG
TalEN2_Xcc CN17   GCAGGCGCTGGAGACGGTGCAGCGGCTGTTGCCGGTGCTGTGCCAGGCCCATGGCCTGACCCCGGAGCAG
TalEN3_Xcc CN18   GCAGGCGCTGGAGACGGTGCAGGCGCTGTTGCCGGTGCTGTGCCAGGCCCATGGCCTGACCCCGCAGCAG


                          2320      2330      2340      2350      2360      2370      2380
                  ....|....|....|....|....|....|....|....|....|....|....|....|....|....|
TalEN5_Xcc XY1-1  GTGGTGGCCATCGCCAGCAATGGCGGCGGCAGGCCGGCGCTGGAGAGCATTGTTGCCCAGTTATCTCGCC
TalEN6_Xcc XY1-2  GTGGTGGCCATCGCCAGCAATGGCGGCGGCAAGCCGGCGCTGGAGAGCATTGTTGCCCAGTTATCTCGCC
TalEN7_Xcc XY2-1  GTGGTGGCCATCGCCAGCAATGGCGGCGGCAGGCCGGCGCTGGAGAGCATTGTTGCCCAGTTATCTCGCC
TalEN8_Xcc XY2-2  GTGGTGGCCATCGCCAGCAATGGCGGCGGCAGGCCGGCGCTGGAGAGCATTGTTGCCCAGTTATCTCGCC
TalEN1_Xcc CN12   GTGGTGGCCATCGCCAGCAATGGCGGCGGCAGGCCGGCGCTGGAGAGCATTGTTGCCCAGTTATCTCGCC
TalEN2_Xcc CN17   GTGGTGGCCATCGCCAGCAATGGCGGCGGCAGGCCGGCGCTGGAGAGCATTGTTGCCCAGTTATCTCGCC
TalEN3_Xcc CN18   GTGGTGGCCATCGCCAGCAATGGCGGCGGCAGGCCGGCGCTGGAGAGCATTGTTGCCCAGTTATCTCGCC

                          2390      2400      2410      2420      2430      2440      2450
                  ....|....|....|....|....|....|....|....|....|....|....|....|....|....|
TalEN5_Xcc XY1-1  CTGATCCGGCGTTGGCCGCGTTGACCAACGACCACCTCGTCGCCTTGGCCTGCCTCGGCGGACGTCCTGC
TalEN6_Xcc XY1-2  CTGATCCGGCGTTGGCCGCGTTGACCAACGACCACCTCGTCGCCTTGGCCTGCCTCGGCGGACGTCCTGC
TalEN7_Xcc XY2-1  CTGATCCGGCGTTGGCCGCGTTGACCAACGACCACCTCGTCGCCTTGGCCTGCCTCGGCGGACGTCCTGC
TalEN8_Xcc XY2-2  CTGATCCGGCGTTGGCCGCGTTGACCAACGACCACCTCGTCGCCTTGGCCTGCCTCGGCGGACGTCCTGC
TalEN1_Xcc CN12   CTGATCCGGCGTTGGCCGCGTTGACCAACGACCACCTCGTCGCCTTGGCCTGCCTCGGCGGACGTCCTGC
TalEN2_Xcc CN17   CTGATCCGGCGTTGGCCGCGTTGACCAACGACCACCTCGTCGCCTTGGCCTGCCTCGGCGGACGTCCTGC
TalEN3_Xcc CN18   CTGATCCGGCGTTGGCCGCGTTGACCAACGACCACCTCGTCGCCTTGGCCTGCCTCGGCGGACGTCCTGC

                          2460      2470      2480      2490      2500      2510      2520
                  ....|....|....|....|....|....|....|....|....|....|....|....|....|....|
TalEN5_Xcc XY1-1  GCTGGATGCAGTGAAAAAGGGATTGCCGCACGCGCCGGCCTTGATCAAAAGAACCAATCGCCGTATTCCC
TalEN6_Xcc XY1-2  GCTGGATGCAGTGAAAAAGGGATTGCCGCACGCGCCGGCCTTGATCAAAAGAACCAATCGCCGTATTCCC
TalEN7_Xcc XY2-1  GCTGGATGCAGTGAAAAAGGGATTGCCGCACGCGCCGGCCTTGATCAAAAGAACCAATCGCCGTATTCCC
TalEN8_Xcc XY2-2  GCTGGATGCAGTGAAAAAGGGATTGCCGCACGCGCCGGCCTTGATCAAAAGAACCAATCGCCGTATTCCC
TalEN1_Xcc CN12   GCTGGATGCAGTGAAAAAGGGATTGCCGCACGCGCCGGCCTTGATCAAAAGAACCAATCGCCGTATTCCC
TalEN2_Xcc CN17   GCTGGATGCAGTGAAAAAGGGATTGCCGCACGCGCCGGCCTTGATCAAAAGAACCAATCGCCGTATTCCC
TalEN3_Xcc CN18   GCTGGATGCAGTGAAAAAGGGATTGCCGCACGCGCCGGCCTTGATCAAAAGAACCAATCGCCGTATTCCC

                          2530      2540      2550      2560      2570      2580      2590
                  ....|....|....|....|....|....|....|....|....|....|....|....|....|....|
TalEN5_Xcc XY1-1  GAACGCACATCCCATCGCGTTGCCGACCACACGCAAGTTGTTCGCGTGCTGGGTTTTTTCCAGTGCCACT
TalEN6_Xcc XY1-2  GAACGCACATCCCATCGCGTTGCCGACCACACGCAAGTTGTTCGCGTGCTGGGTTTTTTCCAGTGCCACT
TalEN7_Xcc XY2-1  GAACGCACATCCCATCGCGTTGCCGACCACACGCAAGTTGTTCGCGTGCTGGGTTTTTTCCAGTGCCACT
TalEN8_Xcc XY2-2  GAACGCACATCCCATCGCGTTGCCGACCACACGCAAGTTGTTCGCGTGCTGGGTTTTTTCCAGTGCCACT
TalEN1_Xcc CN12   GAACGCACATCGCATCTCGTTGCCGACCACACGCAAGTTGTTCGCGTGCTGGGTTTTTTCCAGTGCCACT
TalEN2_Xcc CN17   GAACGCACATCGCATCTCGTTGCCGACCACACGCAAGTTGTTCGCGTGCTGGGTTTTTTCCAGTGCCACT
TalEN3_Xcc CN18   GAACGCACATCCCATCGCGTTGCCGACCACGCGCAAGTGGTTCGCGTGCTGGGTTTTTTCCAGTGCCACT

                          2600      2610      2620      2630      2640      2650      2660
                  ....|....|....|....|....|....|....|....|....|....|....|....|....|....|
TalEN5_Xcc XY1-1  CCCATCCAGCGCAAGCATTTGGTGAAACCATGACGCAGTTCGGGATGAGCAGGCACGGGTTGTTACAGCT
TalEN6_Xcc XY1-2  CCCATCCAGCGCAAGCATTTGATGAAACCATGACGCAGTTCGGGATGAGCAGGCACGGGTTGTTACAGCT
TalEN7_Xcc XY2-1  CCCATCCAGCGCAAGCATTTGATGAAGCCATGACGCAGTTCGGGATGAGCAGGCACGGGTTGTTACAGCT
TalEN8_Xcc XY2-2  CCCATCCAGCGCAAGCATTTGATGAAGCCATGACGCAGTTCGGGATGAGCAGGCACGGGTTGTTACAGCT
TalEN1_Xcc CN12   CCCATCCAGCGCAAGCATTTGATGAAGCCATGACGCAGTTCGGGATGAGCAGGCACGGGTTGTTACAGCT
TalEN2_Xcc CN17   CCCATCCAGCGCAAGCATTTGATGAAGCCATGACGCAGTTCGGGATGAGCAGGCACGGGTTGTTACAGCT
TalEN3_Xcc CN18   CCCACCCAGCGCAAGCATTTGATGACGCCATGACGCAGTTCGGGATGAGCAGGCACGGGTTGTTACAGCT

                          2670      2680      2690      2700      2710      2720      2730
                  ....|....|....|....|....|....|....|....|....|....|....|....|....|....|
TalEN5_Xcc XY1-1  ATTTCGCAGGGTGGGCGTCACCGAACTCGAAGCCCGCAGTGGAACGCTCCCCCCAGCCTCGCAGCGTTGG
TalEN6_Xcc XY1-2  ATTTCGCAGGGTGGGCGTCACCGAACTCGAAGCCCGCAGTGGAACGCTCCCCCCAGCCTCGCAGCGTTGG
TalEN7_Xcc XY2-1  ATTTCGCAGGGTGGGCGTCACCGAACTCGAAGCCCGCAGTGGAACGCTCCCCCCAGCCTCGCAGCGTTGG
TalEN8_Xcc XY2-2  ATTTCGCAGGGTGGGCGTCACCGAACTCGAAGCCCGCAGTGGAACGCTCCCCCCAGCCTCGCAGCGTTGG
TalEN1_Xcc CN12   ATTTCGCAGGGTGGGCGTCACCGAACTCGAAGCCCGCAGTGGAACGCTCCCCCCAGCCTCGCAGCGTTGG
TalEN2_Xcc CN17   ATTTCGCAGGGTGGGCGTCACCGAACTCGAAGCCCGCAGTGGAACGCTCCCCCCAGCCTCGCAGCGTTGG
TalEN3_Xcc CN18   CTTTCGCAGAGTGGGCGTCACCGAACTCGAAGCCCGCAGTGGAACGCTCCCCCCAGCCTCGCAGCGTTGG

                          2740      2750      2760      2770      2780      2790      2800
                  ....|....|....|....|....|....|....|....|....|....|....|....|....|....|
TalEN5_Xcc XY1-1  CACCGTATGCTCCAAGCATCAGGGATGAAAAGGGCCAAACCGTCCTCTGCTTCGGCTCAAACGCCGGACC
TalEN6_Xcc XY1-2  CACCGTATGCTCCAAGCATCAGGGATGAAAAGGGCGAAACCGTCCTCTGCTTCGGCTCAAACGCCGGACC
TalEN7_Xcc XY2-1  CACCGTATGCTCCAAGCATCAGGGATGAAAAGGGCGAAACCGTCCTCTGCTTCGGCTCAAACGCCGGACC
TalEN8_Xcc XY2-2  CACCGTATGCTCCAAGCATCAGGGATGAAAAGGGCCAAACCGTCCTCTGCTTCGGCTCAAACGCCGGACC
TalEN1_Xcc CN12   CACCGTATGCTCCAAGCATCAGGGATGAAAAGGGCGAAACCGTCCTCTGCTTCGGCTCAAACGCCGGACC
TalEN2_Xcc CN17   CACCGTATGCTCCAAGCATCAGGGATGAAAAGGGCGAAACCGTCCTCTGCTTCGGCTCAAACGCCGGACC
TalEN3_Xcc CN18   GACCGTATCCTCCAGGCATCAGGGATGAAAAGGGCCAAACCGTCCCCTACTTCAACTCAAACGCCGGATC

                          2810      2820      2830      2840      2850      2860      2870
                  ....|....|....|....|....|....|....|....|....|....|....|....|....|....|
TalEN5_Xcc XY1-1  AGGAGTCTTTGCATGCATTCGCCGATTCGCTGGAGCGTGAGCTGGATGCGCCCAGCCCAATGGACCAGGC
TalEN6_Xcc XY1-2  AGGAGTCTTTGCATGCATTCGCCGATTCGCTGGAGCGTGAGCTGGATGCGCCCAGCCCAATGGACCAGGC
TalEN7_Xcc XY2-1  AGGAGTCTTTGCATGCATTCGCCGATTCGCTGGAGCGTGAGCTGGATGCGCCCAGCCCAATGGACCAGGC
TalEN8_Xcc XY2-2  AGGAGTCTTTGCATGCATTCGCCGATTCGCTGGAGAGCGAGCTGGATGCGCCCAGCCCAATGGACCAGGC
TalEN1_Xcc CN12   AGGAGTCTTTGCATGCATTCGCCGATTCGCTGGAGCGTGACCTTGATGCGCCTAGCCCAATGCACGAGGG
TalEN2_Xcc CN17   AGGAGTCTTTGCATGCATTCGCCGATTCGCTGGAGCGTGAGCTGGATGCGCCCAGCCCAATGGACCAGGC
TalEN3_Xcc CN18   AGGCGTCTTTGCATGCATTCGCCGATTCGCTGGAGCGTGACCTTGATGCGCCTAGCCCAATGCACGAGGG

                          2880      2890      2900      2910      2920      2930      2940
                  ....|....|....|....|....|....|....|....|....|....|....|....|....|....|
TalEN5_Xcc XY1-1  AGGCCAGGCGCTGGCAAGCAGCCGC---AAACGGTCCCGATCGGATCGGTCTGTCACCGGCTCCTCCGCA
TalEN6_Xcc XY1-2  AGGCCAGGCGCTGGCAAGCAGCCGC---AAACGGTCCCGATCGGATCGTTCTGTCACCGGCTCCTCCGCA
TalEN7_Xcc XY2-1  AGGCCAGGCGCTGGCAAGCAGCCGC---AAACGGTCCCGATCGGATCGTTCTGTCACCGGCTCCTCCGCA
TalEN8_Xcc XY2-2  AGGCCAGGCGCTGGCAAGCAGCCGC---AAACGGTCCCGATCGGATCGTTCTGTCACCGGCTCCTCCGCA
TalEN1_Xcc CN12   AGATCAGACGCGGGCAAGCAGCCGT---AAACGGTCCCGATCGGATCGTGCTGTCACCGGTCCCTCCGCA
TalEN2_Xcc CN17   AGGCCAGGCGCTGGCAAGCAGCCGCCGCAAACGGTCCCGATCGGATCGTTCTGTCACCGGCTCCTCCGCA
TalEN3_Xcc CN18   AGATCAGACGCGGGCAAGCAGCCGCCGCAAACGGTCCCGATCGGATCGTGCTGTCACCGGTCCCTCCGCA

                          2950      2960      2970      2980      2990      3000      3010
                  ....|....|....|....|....|....|....|....|....|....|....|....|....|....|
TalEN5_Xcc XY1-1  CAGCAAGCTGTCGAGGTGCTCGTTCCCGAACAGCGCGATGCGTTGCATTTGCCCCTCAGCTG---GGGTG
TalEN6_Xcc XY1-2  CAGCAAGCTGTCGAGGTGCTCGTTCCCGAACAGCGCGATGCGTTGCATTTGCCCCTCAGCTG---GGGTG
TalEN7_Xcc XY2-1  CAGCAAGCTGTCGAGGTGCTCGTTCCCGAACAGCGCGATGCGTTGCATTTGCCCCTCAGCTG---GGGTG
TalEN8_Xcc XY2-2  CAGCAAGCTGTCGAGGTGCTCGTTCCCGAACAGCGCGATGCGTTGCATTTGCCCCTCAGCTG---GGGTG
TalEN1_Xcc CN12   CAGCAATCGTTCGAGGTGCGCGTTCCCGAACAGCGCGATGCGTTGCATTTGCCCCTCCTCAGCTGGGGTG
TalEN2_Xcc CN17   CAGCAAGCTGTCGAGGTGCTCGTTCCCGAACAGCGCGATGCGTTGCATTTGCCCCTCCTCAGCTGGGGTG
TalEN3_Xcc CN18   CAGCAATCGTTCGAGGTGCGCGTTCCCGAACAGCGCGATGCGTTGCATTTGCCCCTCCTCAGCTGGGGTG

                          3020      3030      3040      3050      3060      3070      3080
                  ....|....|....|....|....|....|....|....|....|....|....|....|....|....|
TalEN5_Xcc XY1-1  TAAAACGCCCGCGTACCAGGATCGGCGGCGGCCTCCTGGATCCTGGTACGCCCATGGATGCCGACCTGGT
TalEN6_Xcc XY1-2  TAAAACGCCCGCGTACCAGGATCGGCGGCGGCCTCCTGGATCCTGGTACGCCCATGGATGCCGACCTGGT
TalEN7_Xcc XY2-1  TAAAACGCCCGCGTACCAGGATCGGCGGCCTCCTCCTGGATCCTGGTACGCCCATGGATGCCGACCTGGT
TalEN8_Xcc XY2-2  TAAAACGCCCGCGTACCAGGATCGGCGGCGGCCTCCTGGATCCTGGTACGCCCATGGATGCCGACCTGGT
TalEN1_Xcc CN12   TAAAACGCCCGCGTACCAGGATCGGCGGC---CTCCTGGATCCTGGTACGCCCATGGATGCCGACCTGGT
TalEN2_Xcc CN17   TAAAACGCCCGCGTACCAGGATCGGCGGC---CTCCTGGATCCTGGTACGCCCATGGATGCCGACCTGGT
TalEN3_Xcc CN18   TAAAACGCCCGCGTACCAGGATCGGCGGC---CTCCTGGATCCTGGTACGCCCATGGATGCCGACCTGGT

                          3090      3100      3110      3120      3130      3140      3150
                  ....|....|....|....|....|....|....|....|....|....|....|....|....|....|
TalEN5_Xcc XY1-1  AGCGTCCAGTACCGTGGTTTGGGAACAAGATGCGGACCCCTTCGCAGGGACAGCGGATGATTTCCCGGCA
TalEN6_Xcc XY1-2  AGCGTCCAGTACCGTGGTTTGGGAACAAGATGCGGACCCCTTCGCAGGGACAGCGGATGATTTCCCGGCA
TalEN7_Xcc XY2-1  AGCGTCCAGTACCGTGGTTTGGGAACAAGATGCGGACCCCTTCGCAGGGACAGCGGATGATTTCCCGGCA
TalEN8_Xcc XY2-2  AGCGTCCAGTACCGTGGTTTGGGAACAAGATGCGGACCCCTTCGCAGGGACAGCGGATGATTTCCCGGCA
TalEN1_Xcc CN12   AGCGTCCAGCACCGTGGTTTGGGAACAAGATGCGGACCCCTTCGCAGGGACAGCGGATGATTTCCCGGCA
TalEN2_Xcc CN17   AGCGTCCAGCACCGTGGTTTGGGAACAAGATGCGGACCCCTTCGCAGGGACAGCGGATGATTTCCCGGCA
TalEN3_Xcc CN18   AGCGTCCAGCACCGTGGTTTGGGAACAAGATGCGGACCCCTTCGCAGGGACAGCGGATGATTTCCCGGCA

                          3160      3170      3180      3190
                  ....|....|....|....|....|....|....|....|....|...
TalEN5_Xcc XY1-1  TTCAACGAAGAGGAGCTCGCATGGTTGATGGAGATATTGCCTCAGTGA
TalEN6_Xcc XY1-2  TTCAACGAAGAGGAGCTCGCATGGTTGATGGAGAGATTGCCTCAGTGA
TalEN7_Xcc XY2-1  TTCAACGAAGAGGAGCTCGCATGGTTGATGGAGAGATTGCCTCAGTGA
TalEN8_Xcc XY2-2  TTCAACGAAGAGGAGCTCGCATGGTTGATGGAGATATTGCCTCAGTGA
TalEN1_Xcc CN12   TTCAACGAAGAGGAGCTCGCATGGTTGATGGAGCTATTGCCTCATTGA
TalEN2_Xcc CN17   TTCAACGAAGAGGAGCTCGCATGGTTGATGGAGCTATTGCCTCATTGA
TalEN3_Xcc CN18   TTCAACGAAGAGGAGCTCGCATGGTTGATGGAGCTATTGCCTCATTGA
```

**Figure S1**. Alignment of coding regions of the TALE genes from strains XY1-1, XY1-2, XY2-1, and XY2-2 (this work) and CN12, CN17, and CN18 (Denancé et al., 2018). TALEs from CN12, CN17, CN18 belong to the Tal15g group according to Denancé et al. (2018). The TALE gene names are in accordance with the naming of the AnnoTale suite. Repeats in the central part of genes are coloured.

**Table S1**. Primers used for RT-PCR.

| Target gene | Accession Number | Sequence | Note |
| --- | --- | --- | --- |
| ERF121 (target) | XM_013739306 | F 5'- CCCTGTTTCACAACCATCTTAC -3'  R 5'- GAGGCGGAAGAACTTCACTA -3' | - |
| Actin-2 (reference) | XM_013731369.1 | F 5'- CTGAGGAGCACCCGGTTCTTCTTA -3'  R 5'- CGGAGGATGGCGTGTGGTAGA -3' | Both primers from *Afrin, K. S., Rahim, M. A., Park, J. I., Natarajan, S., Kim, H. T., & Nou, I. S. (2018). Identification of NBS-encoding genes linked to black rot resistance in cabbage (Brassica oleracea var. capitata). Molecular biology reports, 45(5), 773-785.* |
| GAPDH (reference) | JN571725.1 | F 5'- GAAAGGTGCTTCCACAGCTC -3'  R 5'- GTCGCAGCTTTCTCGAGTCT -3' | Both primers from *Villeth, G. R., Carmo, L. S., Silva, L. P., Santos, M. F., de Oliveira Neto, O. B., Grossi-de-Sá, M. F., ... & Mehta, A. (2016). Identification of proteins in susceptible and resistant Brassica oleracea responsive to Xanthomonas campestris pv. campestris infection. Journal of proteomics, 143, 278-285.* |
| EF1α (reference) | AT5G60390 | F 5'- TGAGCATGCTCTTCTTGCTTTCA -3'  R 5'- TACCTAGCCTTGGAGTACTTAGG -3' | Forward primer with a single-nucleotide modification from *Santos, C., Nogueira, F., Domont, G. B., Fontes, W., Prado, G. S., Habibi, P., ... & Franco, O. L. (2019). Proteomic analysis and functional validation of a Brassica oleracea endochitinase involved in resistance to Xanthomonas campestris. Frontiers in plant science, 10, 414.* |

**Table S2**. Similarity between aa sequences of the different TALE-activated ERFs.

| aa identity/similarity, % (blosum62 mt2 alignment matrix, AlignX application) | XP_013594760^1^  *B.oleracea*  ERF#121(phylogenetic group Xb-L**)* | LOC_Os07g47790.1^2^  *O. sativa*  ERF#067(phylogenetic group VIIa)* | Os09g0571700^3,4^  *O. sativa*  ERF#123(phylogenetic group IXc)* | VAH18767.1^5^  *T. turgidum*  TaERF_1BL (phylogenetic group Ia***)* |
| --- | --- | --- | --- | --- |
| XP_013594760  *B. oleracea*  ERF#121 | - | 12,5/23 | 9,6/18,5 | 5,1/9,3 |
| LOC_Os07g47790.1  *O. sativa*  ERF#067 | 12,5/23 | - | 22,1/29,5 | 9,3/18,3 |
| Os09g0571700  *O. sativa*  ERF#123 | 9,6/18,5 | 22,1/29,5 | - | 5,8/10,9 |
| Traes_1BL_F0C52C5DF  *T. turgidum*  TaERF_1BL | 5,1/9,3 | 9,3/18,3 | 5,8/10,9 | - |

* Nakano T, Suzuki K, Fujimura T, Shinshi H. 2006. Genome-wide analysis of the ERF gene family in Arabidopsis and rice. *Plant physiology*, **140**: 411-432.

** Based on similarity with Arabidopsis proteins

*** Based on similarity with rice proteins

^1^this work

^2^Pérez-Quintero AL, Rodriguez-R LM, Dereeper A, López C, Koebnik R, Szurek B, Cunnac S. 2013. An improved method for TAL effectors DNA-binding sites prediction reveals functional convergence in TAL repertoires of *Xanthomonas oryzae* strains. *PloS one* **8**: e68464.

^3^Tran TT, Pérez-Quintero AL, Wonni I, Carpenter SC, Yu Y, Wang L, leach JE, Verdier V, Cunnac S, Bogdanove AJ et al. 2018. Functional analysis of African *Xanthomonas oryzae* pv. *oryzae* TALomes reveals a new susceptibility gene in bacterial leaf blight of rice. *PLoS pathogens* **14**: e1007092.

^4^Wang L, Rinaldi FC, Singh P, Doyle EL, Dubrow ZE, Tran TT, Pérez-Quintero AL, Szurek B, Bogdanove AJ. 2017. TAL effectors drive transcription bidirectionally in plants. *Molecular plant* **10**: 285-296.

5 Peng Z, Hu Y, Zhang J, Huguet-Tapia JC, Block AK, Park S, Sapkota S., Liu Z, Liu S, White FF. 2019. *Xanthomonas translucens* commandeers the host rate-limiting step in ABA biosynthesis for disease susceptibility. *Proceedings of the National Academy of Sciences* **116**: 20938-20946.
